# CATD: A reproducible pipeline for selecting cell-type deconvolution methods across tissues

**DOI:** 10.1101/2023.01.19.523443

**Authors:** Anna Vathrakokoili Pournara, Zhichao Miao, Ozgur Yilimaz Beker, Nadja Nolte, Alvis Brazma, Irene Papatheodorou

**Affiliations:** European Molecular Biology Laboratory, European Bioinformatics Institute, EMBL-EBI, Wellcome Genome Campus, Cambridgeshire, CB10 1SD, UK; Open Targets, Wellcome Genome Campus, Hixton, Cambridgeshire CB10 1SD, UK; GMU-GIBH Joint School of Life Sciences, Guangzhou Laboratory, Guangzhou Medical University, Guangzhou, China; Faculty of Engineering and Natural Sciences, Sabanci University, Tuzla 34956, Istanbul, Turkey; Department of Biotechnology and Systems Biology, National Institute of Biology, Ljubljana, Slovenia

## Abstract

Cell-type deconvolution methods aim to infer cell composition from bulk transcriptomic data. The proliferation of developed methods, coupled with inconsistent results obtained in many cases, highlights the pressing need for guidance in the selection of appropriate methods. Additionally, the growing accessibility of single-cell RNA sequencing datasets, often accompanied by bulk expression from related samples, enable the benchmark of existing methods. In this study, we conduct a comprehensive assessment of 31 methods, utilising single-cell RNA-sequencing data from diverse human and mouse tissues. Employing various simulation scenarios, we reveal the efficacy of regression-based deconvolution methods, highlighting their sensitivity to reference choices. We investigate the impact of bulk-reference differences, incorporating variables such as sample, study and technology. We provide validation using a gold standard dataset from mononuclear cells and suggest a consensus prediction of proportions when ground truth is not available. We validated the consensus method on data from the stomach and studied its spillover effect. Importantly, we propose the use of the Critical Assessment of Transcriptomic Deconvolution (CATD) pipeline which encompasses functionalities for generating references and pseudo-bulks and running implemented deconvolution methods. CATD streamlines simultaneous deconvolution of numerous bulk samples, providing a practical solution for speeding up the evaluation of newly developed methods.

**Availability:** https://github.com/Papatheodorou-Group/CATD_snakemake

## 1. Introduction

There is a growing interest in understanding the level of heterogeneity and the importance of cell-type abundances in healthy and diseased tissues. Transcriptomic heterogeneity and cell-type composition reveal critical features of tissue functionality. For example, in cancer, immune cells can either be recruited in the vicinity of the tumour to contribute to cancer progression, or they can play a tumour-suppressive role by recognising and killing abnormal cells (Hanahan and Coussens, 2012; Dumont *et al*., 2013; Taube *et al*., 2018; Jorge *et al*., 2020). High infiltration from specific immune cells can be predictive of the disease outcome, stage and aid the treatment selection (Zhang *et al*., 2013). However, determining the composition of a tissue sample can present significant challenges. While single-cell RNA sequencing techniques offer valuable insights into tissue heterogeneity, challenges such as technical handling, sorting of specific cell types, sub-optimal dissociation and tissue vulnerability may affect the accurate determination of cell type proportions in single-cell data (Denisenko *et al*., 2020). Furthermore, conventional methods for quantifying cell abundances, such as flow cytometry and immunohistochemistry, rely on prior knowledge of cell markers, depend on interpretation bias and are not easily scalable (Matos *et al*., 2010; Patrick *et al*., 2020). Additionally, there are only a handful of this type of datasets available and mostly deriving from non-solid tissues such as blood (Finotello *et al*., 2019; Monaco *et al*., 2019). Taking all the above into account, computational deconvolution methods show a great advantage, since cell proportions from hundreds of samples can be obtained computationally very fast – in a matter of seconds to a few hours depending on the selected method. As a result, cell-type deconvolution methods –also called decomposition methods– provide a cost-effective way of measuring cell proportions when compared to experimental methods. Moreover, deconvolution methods can be very useful for inferring cell type compositions from historical bulk RNA-seq data from large databases such as GTEx (GTEx Consortium, 2017), TCGA (The International Cancer Genome Consortium, 2010), TARGET (https://ocg.cancer.gov/programs/target) and GEO (Edgar, 2002) in which it would be impossible to repeat experiments on.

Cell-type deconvolution can contribute significantly in answering critical biological questions and understanding disease mechanisms to a greater extent. Specifically, it can unravel how cell proportions can affect certain phenotypes or specific clinical features available within genomic databases (Donovan *et al*., 2020a). Additionally it can serve as a powerful tool for mitigating the confounding effect of cell-type proportions in bulk differential expression analysis and as a result help in the identification of reliable biomarkers for diseases and perturbed processes and pathways with cell-type resolution (Inkeles *et al*., 2016). Moreover, in the context of Transcriptome-wide association studies (TWAS) deconvolution can rectify cell abundance bias in the expression quantitative trait loci (eQTL) analysis and facilitate the identification of cell-type specific eQTLs (Lowe and Rakyan, 2014). Finally, the abundance of bulk RNA-seq samples present a unique opportunity for training machine learning models tailored to phenotype prediction, a task that is often challenging with single-cell data due to limited sample sizes. Importantly, by integrating cell composition, we effectively enrich the information content that bulk samples are missing due to the experimental method principles.

Deconvolution methodologies can be formally categorised into three main groups based on their input requirements: supervised (reference-based), semi-supervised, and unsupervised (complete). Supervised deconvolution methods necessitate an expression matrix (typically from bulk RNA-sequencing or microarray data) to be deconvolved as well as cell-type-specific information. This specific information can be derived from sources like flow cytometry, single-cell RNA-seq data, or marker gene lists. The above supervised methods can further be divided into two subcategories: bulk when the reference is either a signature of sorted cell types or a marker gene list and single-cell when the reference is a single-cell dataset. Well-known ‘bulk’ deconvolution methods include CIBERSORT (absolute) (Newman *et al*., 2015), FARDEEP (Hao *et al*., 2019), NNLS (non-negative least squares) and EpiDISH (Teschendorff *et al*., 2017)). Concurrently, there are contemporaneous deconvolution methods that harness available single-cell datasets to extract cell type-specific features and subsequently perform deconvolution. Examples of such methods include Bseq-sc (Baron, Veres, Samuel L. Wolock, *et al*., 2016), DWLS (Tsoucas *et al*., 2019) and MuSiC (Wang *et al*., 2019a) and others (Frishberg *et al*., 2019; Jew *et al*., 2020; Dong *et al*., 2021). These approaches incorporate feature selection methods to choose informative genes for deconvolution, such as DWLS-MAST, MuSIC-informative genes, and AutogeneS. All the above methods output cell proportions, enabling both intra-sample and inter-sample comparisons. In the supervised category, gene set enrichment methods such as xCell (Aran, Hu and Butte, 2017), MCP-counter (Becht *et al*., 2016), and SaVant (Lopez *et al*., 2017) can also be included. However, these methods use an enrichment-based approach and output enrichment scores, not percentages. These scores can be compared across samples, but do not allow intra-sample comparison. Unsupervised methods, on the other hand, such as deconf (Repsilber *et al*., 2010) and CDSeq (Kang *et al*., 2019) require only a bulk sample and the number of components (representing the cell types to be identified) as input. These methods estimate both the cell-type-specific expression and cell fractions simultaneously. Semi-supervised methods, such as ssKL and ssFrobenius, can also estimate both the cell-specific expression matrix and the matrix of cell proportions at the same time. They utilise a list of pre-determined markers, the expected number of components, and the bulk matrix as input. These semi-supervised methods employ a block-descent approach to estimate both matrices (Gaujoux and Seoighe, 2012). In this study, our primary focus is on comparing supervised and semi-supervised methods, and we do not extensively explore unsupervised (complete) deconvolution methods. Unsupervised methods are particularly valuable when the cell composition is entirely unknown or cell signatures for the cell types of interest are not available. These methods have been thoroughly investigated in previous benchmarks to assess their performance and underlie their potential (Jaakkola and Elo, 2021; Jin and Liu, 2021; Sutton *et al*., 2022). Lastly, there is a separate class of methods known as ‘expression deconvolution methods’. These methods tackle the inverse problem; they aim to find the cell type-specific expression matrix given prior knowledge of cell proportions and bulk expression data. Methods falling under this category include Rodeo (Jaakkola and Elo, 2021), csSAM (Shen-Orr *et al*., 2010), Deblender (Dimitrakopoulou *et al*., 2018), cs-lsfit (Gaujoux and Seoighe, 2012), and cs-prog (Gong *et al*., 2011). Previous benchmark studies have evaluated the performance of these methods, but due to their distinct design and research focus, we have not included those in our study.

The field of deconvolution methods has witnessed significant growth in recent years, driven by the complex nature of the deconvolution problem and the ongoing development of new and more sophisticated techniques (Menden *et al*., 2020). Over the past decade, there have been notable efforts to independently evaluate deconvolution methods (Avila Cobos *et al*., 2020a; Jin and Liu, 2021). These efforts explore various aspects, such as the effectiveness of simulations, method performance, the importance of reference data, the impact of data normalisation, the influence of different technologies, and the granularity of cell-type information in the data (Teschendorff *et al*., 2017; Jiménez-Sánchez, Cast and Miller, 2019; Sturm *et al*., 2019; Nadel *et al*., 2021; Sutton *et al*., 2022; White *et al*., 2022; Alonso-Moreda *et al*., 2023; Cobos *et al*., 2023). However, as our knowledge in this field continues to expand, and new methods are introduced each year, there is a growing need for comprehensive, reproducible, and versatile benchmarking pipelines that can accommodate both existing and newly developed methods(Garmire *et al*., 2024). This need is especially pressing because different deconvolution methods may have varying input formats, design methodologies, preprocessing steps, and output results. Therefore, a centralised and efficient platform is essential to streamline the evaluation process, facilitating a quicker and more unified assessment of deconvolution methods. Furthermore, while previous studies have primarily focused on deconvolution in peripheral blood mononuclear cells (PBMC) samples or in a limited number of tissues each time, there is a growing demand for research that covers more tissues and datasets as more and more cell type-specific information becomes available(Maden *et al*., 2023).

To address these challenges, we have designed a robust and reproducible Snakemake pipeline for evaluating thirty-one available deconvolution methods and obtaining results on cell abundances from real data. Our evaluation involves nineteen single-cell datasets from diverse sources, covering five different tissues with multiple datasets per tissue, enabling us to generate a wide range of simulated and reference data for method evaluation. Our pipeline has also incorporated various simulation approaches, leveraging single-cell datasets and suggesting parameter settings to generate realistic simulated data across different tissue types. Additionally, we study the impact of preprocessing methods and the effect of distinct technologies in the reference data. Deconvolution performance is finally evaluated on real data from PBMC samples with ground truth. Moreover, we introduced a consensus deconvolution approach by combining well-performing methods and validated the results on stomach samples from GTEx using abundances from tissue slides as our ground truth. Finally, we assess the spillover effect of the consensus method on purified samples from cerebral cortex and lung tissue. One notable feature of our pipeline is its adaptability to newly developed deconvolution methods that require independent benchmarking. Furthermore, researchers can employ our pipeline to simultaneously deconvolve multiple biological samples, obtaining results from implemented methods and benefiting from the pipeline’s visualisations. By assessing the consistency of results across methods and references, researchers can subsequently choose the most suitable setup to address their specific research questions.

## 2. Methods

### 2.1. Data Collection for benchmarking

We collected single-cell data from various sources to generate references and simulate bulk data for in-silico experiments. 12 single-cell datasets were collected from human and mouse brain tissues, including dental gyrus, anterolateral motor cortex (ALM), primary visual cortex (ViSP), cerebral cortex and pons, retinas, hippocampus CA1, somatosensory cortex S1, primary motor cortex, cortex areas and glioblastoma cancer tissue. We also collected two single-cell datasets specifically focused on human pancreatic islets to investigate sample effects in the deconvolution process. To understand the impact of different platforms/single-cell technologies in deconvolution we curated three pairs of 10X and Smart-seq2 datasets, with each pair originating from the same study. For this task we covered three tissues: placenta, brain and lung. A comprehensive breakdown of the datasets is provided in Table S1, offering details on the number of samples, cells, genes and cell-types per dataset, covered conditions, and corresponding GEO accession numbers.

### 2.2. Single-cell data (re)analysis

Raw counts and metadata (minimum metadata required: cell-type annotations, cellIDs, sampleIDs) were obtained for each single-cell dataset used for the in-silico benchmarking experiments. Single-cell datasets used for reference generation were re-analysed utilising the SCANPY toolkit (Wolf, Angerer and Theis, 2018). Cells with less than 200 genes and genes expressed in less than 3 cells were filtered out. Cells with more than >5% of mitochondrial genes or too many total counts (dataset-specific threshold) were also removed. To inspect the cell annotation quality all datasets were normalised (total counts = 10e4) and log-transformed, the raw count dimension was kept intact to be used as a deconvolution input in the pipeline. Next, highly variable genes were identified and we performed Principal Component Analysis (PCA), computed the neighbourhood graph as well as performed leiden clustering using different resolution parameters (0.25, 0.5, 1). After clustering, we plotted the data on UMAP space using as colour key: cellType and leiden clusters in order to evaluate the author’s annotations. Cell types annotated from the authors in the original papers as ‘others’, ‘unknown’, ‘low quality’, ‘unclassified’ were removed from the data. Additionally, cell type nomenclature was harmonised for consistency across datasets. This step is particularly useful for the cross-reference and real bulk deconvolution tasks. To validate cell type annotations, cell types were examined using known marker genes. This step involved visualising the expression of genes known to be specific to particular cell types on UMAP space.

### 2.3. Real data validation

To validate the results from benchmarking on real data we collected real bulk samples from various tissues and designed validation experiments using diverse ground truth data. Collection, curation and design of validation experiments are described below. More details can be found in Table S4.

#### Validation on human PBMC samples

We collected bulk RNA-seq data from 9 PBMC human samples. For these samples we have also coupled flow cytometry measurements for the cell type proportions (GSE107572) (Finotello *et al*., 2019). To construct the references for deconvolution we utilised a publicly available single-cell dataset from 14 PBMC samples (GSE150728) (Wilk *et al*., 2020). We harmonised the cell type annotation between the single-cell and the bulk data, ensuring seamless deconvolution analysis and evaluation (Supplementary Figure 8).

#### Validations on GTEx stomach data

We acquired bulk expression data (TPM-normalised) from GTEx portal v8 and focused on the stomach tissue data. Among the 939 stomach samples, 339 were accompanied by histology slides, enabling the calculation of ground truth proportions through QuPath (Bankhead *et al*., 2017) analysis (see Methods 2.12). To use a reliable reference for this task we used the stomach subset of the Human Cell Landscapes single-cell dataset, downloaded from cellxgene portal (CZI Single-Cell Biology Program *et al*., 2023). For a standardised reduction of cell type label granularity we employed an automated ontology-based method using the scOntoMatch (v 0.1.0) package (Song *et al*., 2023).

The adapted code for this task is available at: https://github.com/nadjano/reference_preperation_for_GTEX_deconvolution

#### Deconvolving purified flow cytometry data

To study the spillover effect previously documented in deconvolution we collected and deconvolved bulk samples of purified cell types from two tissue sources, human fetal lung (E-MTAB-9372) and mouse brain (E-GEOD-52564). The data were downloaded from Expression Atlas (Moreno *et al*., 2022). As a lung specific deconvolution reference, we used the subset from the GTEx v7 single-cell dataset. Cell type labels granularity was reduced as described in the previous section. For the deconvolution of mouse brain data, Tabula Muris dataset (The Tabula Sapiens Consortium* *et al*., 2022) (E-ENAD-15) was utilised. To expedite the deconvolution computation we downsampled the above single-cell references. Specifically, for cell types that possessed more than 300 cells, we employed a random sampling approach, selecting 300 cells per cell type. The detailed code implementation for this procedure can be found here: (https://github.com/nadjano/reference_preperation_for_GTEX_deconvolution).

### 2.4. Overview of CATD Snakemake Pipeline

The CATD pipeline is a benchmarking pipeline meant to facilitate the assessment of cell-type deconvolution methods (currently 31) across different simulation scenarios in a standardised way. It also allows the deconvolution of real bulk samples with various input parameters allowing users to deconvolute their own in-house data following our proposed guidelines. The pipeline includes:

- Pseudo-bulk generation methods (from single-cell RNA-seq data) that allow to create diverse pseudo-bulk samples and compare deconvolution methods across different scenarios.
- 17 normalisation methods implemented in the pipeline for the normalisation of the input single-cell reference and the (pseudo) bulk samples
- 4 transformation methods
- 9 Differential Expression tests for the selection of marker genes from single-cell reference data (Seurat, FindMarkers)
- 31 deconvolution methods
- 9 metrics to assess the results when we test deconvolution methods on pseudo-bulks or when ground truth proportions from real data are available.
- A Consensus approach, combining three well-performing deconvolution methods

Moreover the pipeline provides:

- Visualisation of the performance and scalability metrics across methods.
- Visualisation of the consensus deconvolution results (bar plots with cell type proportions per sample)
- Visualisation of the correlation of the results across methods for a given dataset when ground truth is not available.

### 2.5. Generation of pseudo-bulk (synthetic) samples

To benchmark deconvolution methods we designed various pseudo-bulk simulation data using different sampling methodologies. Each simulation starts with splitting the single cell in half (50% testing data and 50% training data). Testing data will be then used to generate the pseudo-bulks. All the simulations require as an input to set the number of samples (n) to be generated and the number of cells to sample from the single cell for each pseudo-bulk (c). Two modes of pseudo-bulk generation have been designed for the purpose of our benchmark. For our simplest simulation (mode 1), the sampling will randomly pick c number of cells with replacement from the single-cell data to generate the first sample, this will be repeated n times until we obtain the final pseudo-bulk dataset. For mode 2, we have designed a more realistic pseudo-bulk generation method which takes into account variability of cell type proportions and cell type heterogeneity for each sample. Hence for mode 2 the generation of pseudo-bulks will require two additional parameters, namely ‘proportional variance’ (*p*) and second, a logical parameter called ‘sampleCT’ that defines whether or not some cell types will be left out during generation. When including all cell types in our pseudo-bulks, if *p* is set to a negative number, then random proportions ranging between 0.01 to 0.99 and summing to 1, will be generated using a uniform distribution. If *p* is positive, then integers between *1* and *1+2p* will be generated using a uniform distribution and then normalised to sum to 1. This enables more precise control of the noise, with the former (1-99 case) being the edge case of the latter (positive *p*) when *p* tends to infinity. If cell types are sampled, a bimodal normal distribution with uniform mode probability is used to achieve the effect and the *p* instead becomes the standard distribution of the kept cell types. In this case, the provided *p* should be positive and ideally large. The absolute value of generated numbers is considered to ensure non-negative proportions and then proportions are normalised to sum to one.

The generated proportions in mode 2 are then scaled by *c* to determine how many cells will be chosen with replacement per cell type from the reference data to generate one sample, which is again repeated *n* times.

### 2.6. Building reference data from single-cell data

To build references used by the deconvolution methods we start by obtaining a single cell RNA-sequencing expression matrix. Different methods require different types of reference input.

- First we have ‘single-cell’ methods that require a single-cell matrix as input. Within the pipeline, this matrix is encoded as ***C0. C0*** is a generic single-cell matrix, with rows being genes and columns being cellIDs, this matrix is used as an input in single-cell deconvolution methods (e.g MuSiC, DWLS, Bisque and others). Single-cell methods also need an additional metadata object encoded as ***phenData*** in our pipeline which is associated with ***C0*** and provides additional information (sampleIDs, cellIDs, cell type labels).
- Next we have reference-based deconvolution methods that require input expression data from purified cell types (e.g FACS data). For these methods we generate the **C1 matrix** which is the averaged (arithmetic mean) expression matrix by cell types, with rows being genes and columns being cell types.
- For marker-based methods we create the ***C2* reference**. C2 is a list of marker genes obtained from differential gene expression analysis on the single-cell data through the Seurat package. Specifically a combination of *NormalizeData()* and *FindAllMarkers()* functions are being used. Methods that use C2 reference as input include: EPIC, DSA, ssFrobenious, ssKL, deCAMmarker. By default we use the ‘wilcox’ test (Wilcoxon Rank Sum test) to identify marker genes. Moreover we have implemented in the pipeline an additional functionality to use two tests each time and obtain the overlap of marker genes from the two tests. However, this could result in issues while running the pipeline in cases where the overlap of genes is very low.
- Finally, ***refVar*** is the row standard deviations matrix by cell types, similar in structure to ***C1***. refVar is used only by EPIC deconvolution method due to its unique design.

### 2.7. Preprocessing methods

To comprehensively examine the impact of variable pre-processing on both (pseudo)bulk and single-cell matrices, we employed an array of normalisation and transformation methods. While many of the normalisation techniques applied in this study are suitable for both data types, we also incorporated four methods tailored specifically for single-cell data: SCTransform, scran, scatter and Linnorm. A detailed summary of all the methods, including their characteristics, can be found in Table S2.

Furthermore, we investigated the data transformation effects. To explore this aspect, we implemented four distinct transformations: logarithmic (log), square (sqrt), variance-stabilising transformation (vst), or no transformation (linear scale). Finally, we systematically examined the impact of the order of preprocessing steps, considering whether transformation or normalisation should be conducted first. Detailed insights into the implementation of all preprocessing steps and methods are available within the pipeline.

### 2.8. Selection of deconvolution methods

In our benchmarking study, we curated deconvolution methods from existing literature and previous benchmark studies (Avila Cobos *et al*., 2020a; Jaakkola and Elo, 2021; Jin and Liu, 2021; Sutton *et al*., 2022). The selected methods represent diverse categories, including 27 supervised methods, which further break down into 10 single-cell-based methods, 12 reference-based methods, and 4 marker-based methods. Additionally, we incorporated 2 semi-supervised methods (ssKL and ssFrobenious) and 2 unsupervised methods (deconf and CDSeq).

A comprehensive summary of the enlisted methods is available in Table S3, providing details on their characteristics, algorithmic principles, implementation specifics, tool version numbers, and original publications.

### 2.9. Metrics for performance evaluation

To assess the efficacy of deconvolution methods, we conducted a thorough evaluation by comparing the proportions derived by these methods against known ground truth proportions. Depending on the specific task, the ground truth can either be experimentally calculated proportions or proportions derived from simulations. Our pipeline utilises key metrics to evaluate performance, including Pearson correlation coefficients, Spearman correlation coefficients, Root Mean Square Error (RMSE), and weighted RMSE. For weighted RMSE each cell type is assigned a weight based on its proportion value in a way where rare cell types are given bigger weight. Additionally, we incorporated supplementary metrics such as Mean Absolute Error (MAE)/Mean Absolute Deviation (MAD), Euclidean distance, Distance correlation, Cosine Similarity (cos), and R-squared. The notation and equations for calculating these metrics are detailed in the Supplementary Materials. All metrics were computed in R, and the corresponding functions can be accessed at: https://github.com/Functional-Genomics/CATD_snakemake/blob/main/Modules/Res_explore/Res_explore.R.

### 2.10. Metrics for scalability evaluation

In the context of deconvolution, assessing scalability is crucial, particularly when considering the application of these methods to large-scale datasets such as extensive databases with historical bulk data. Evaluating scalability ensures that the computational efficiency of deconvolution can meet the demands of processing substantial volumes of data. Every step of the pipeline is benchmarked automatically by Snakemake’s built-in ‘benchmark’ directive, and the results from these benchmarks are also visualised at the end of the pipeline. Four of the parameters are visualised for initial scalability assessment purposes, namely the time (s), memory (maximum resident set size), mean load (CPU usage divided by total time) and CPU time.

### 2.11. Consensus approach

For the consensus approach we selected three deconvolution methods: DWLS, EpiDISH, FARDEEP. These methods were performing well across our study. The main criteria for inclusion were:

1. well-performing methods overall
2. open-source code,
3. reasonable time of running,
4. stable and they produce results (sometimes methods won’t converge and produce NAs.
5. flexible (e.g they can take as input different references every time to answer biological questions)

The displayed cell type proportions of GTEx stomach, mouse purified brain, and human fetal lung cell types are computed as the averages of deconvolution outcomes obtained through DWLS, EpiDISH, and FARDEEP, per sample and cell type. We also include the mean correlation between the results from the three methods across all samples to show the concordance of results across the methods. Standard deviation (sd) per cell type for each sample across methods is also computed. For the consensus visualisation of deconvolution proportions in GTEx stomach samples, we employed the CiberBarFrazer() function (Donovan *et al*., 2020b). Code available at https://github.com/mkrdonovan/gtex_deconvolution, R version 4.3.1). The resulting proportions were arranged in descending order of the most abundant cell types.

### 2.12. QuPath analysis

To validate our deconvolution results for selected stomach samples, we leveraged histology images downloaded from the GTEx Portal Histology Viewer (https://gtexportal.org/home/histologyPage). Histology files for 339 samples were downloaded, and for analysis, we employed QuPath software (v 0.4.1 on Mac OS X - 11.6.2). A pixel classifier was trained on six slides, allowing us to estimate the areas of background, muscle, mucosa, and submucosal layers. Subsequently, using the assigned areas, we calculated the relative proportions of the three tissue layers on all slides.

For the mapping of relevant cell types to stomach layers, we drew upon insights from the literature on the organisational structure of stomach tissues. Specifically the mapping was done as follows:

- Mucosa: Endocrine cells, enterocytes, goblet cells, peptic cells (parietal cells), and epithelial cells.
- Submucosa: Dendritic cells, endothelial cells, erythroid lineage cells, fibroblasts, macrophages, plasma cells, and stromal cells.
- Muscle: Mesenchymal stem cells, neurons, and smooth muscle cells.

## 3. Results

### 3.1 Computational framework for large-scale analysis of deconvolution (CATD)

We developed a computational framework, called CATD, that enables us to systematically investigate various factors that influence the deconvolution process, in order to provide clear guidelines for the decomposition of bulk tissue expression data. (Figure 1). To benchmark the 31 available deconvolution methods, we collected scRNA-sequencing published datasets from different human and mouse organs (Table S1). In our pipeline, each scRNA-seq dataset is split in half into a training dataset, used as a “reference input” for deconvolution, and a testing dataset that is utilised to generate pseudo-bulk mixtures to be deconvolved afterwards (<u>Methods</u>). The use of pseudo-bulk mixtures allows us to obtain ground truth against which we can evaluate the deconvolution methods. The collected scRNA-seq experiments were generated by numerous single-cell protocols including Smart-Seq2 (SS2) (Picelli *et al*., 2013), 10X (Zheng *et al*., 2017), CEL-seq (Hashimshony *et al*., 2012), Seq-well (Gierahn *et al*., 2017), Drop-Seq (Macosko *et al*., 2015), inDrop (Zilionis *et al*., 2017) and originate from different organs including brain, pancreas, lung, blood (PBMC) and placenta. Moreover, datasets of different granularity were included in the benchmark to account for the effect of cell-type numbers in deconvolution (Table S1). Having included multiple datasets from various sources, the generated pseudo-bulks encompass a wide range of biological environments. Additionally, pseudo-bulk data designed to encompass different levels of noise in order to mimic real bulk samples. In addition, 11 different normalisation and 2 transformation methods (<u>Methods</u>, Table S2) were tested on both pseudo-bulk and reference matrices to select the most suitable combination of preprocessing to be performed prior to deconvolution. After the preprocessing step, we evaluated the performance of the 31 collected deconvolution methods (Table S3). To compare the performance of deconvolution methods we implemented various evaluation metrics in the CATD pipeline. Pearson correlation coefficient (r/pcor), root mean square error (RMSE), average cosine similarity (avgcos), weighted RMSE and Spearman’s correlation are some of the key metrics implemented and used across this study to evaluate the performance of the methods in different scenarios (<u>Methods</u>). We select the best performing methods taking into consideration the methods that have both low RMSE and high R values consistently across different simulation scenarios, datasets and tissues. All the steps of the benchmarking study have been modularised in a snakemake pipeline for systematic exploration and reproducibility of the benchmarking results. The pipeline also allows the evaluation of the scalability of each benchmarking module using additional metrics (<u>Methods</u>). Finally, CATD provides a practical, user-friendly framework to accurately deconvolve real bulk samples with or without ground truth by suggesting a new approach to obtain a fair consensus of the best-performing methods and providing visualisation of the deconvolution results.

**Figure 1.**
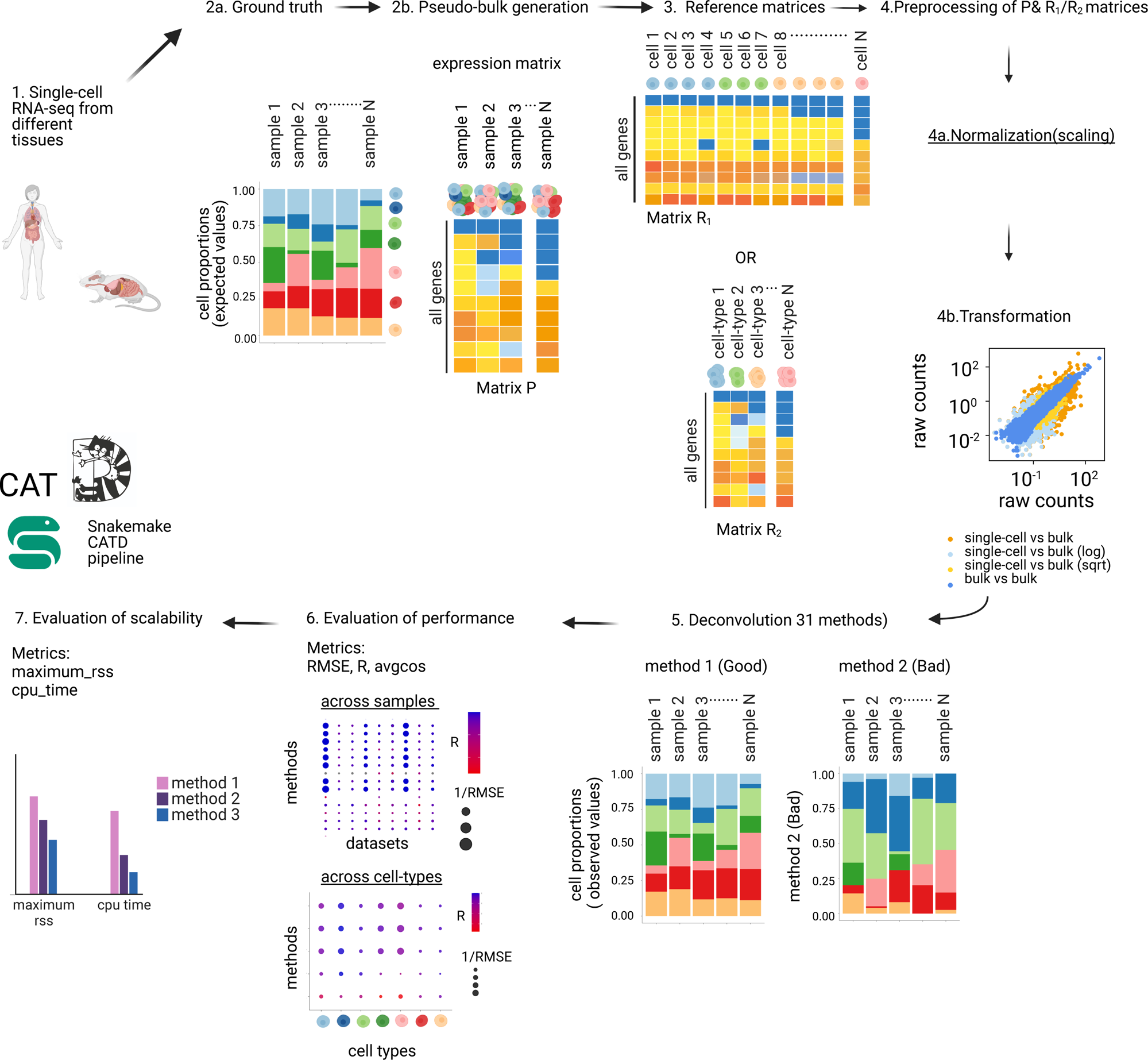
Overview of Critical Assessment of Transcriptomic Deconvolution(CATD) pipeline. Schematic representation of the large-scale benchmarking pipeline. 31 deconvolution methods were evaluated on simulated bulk data generated from scRNA-seq datasets and 4 separate simulation strategies. Furthermore, we test different normalisation and transformation methods on the bulk and reference matrices and we assess the results utilising various evaluation metrics. Finally, we evaluate the scalability of the methods. Every step of the benchmarking has been modularised using snakemake.

### 3.2 Realistic pseudo-bulk simulations for systematic evaluation and method development

#### Objective

Efforts to benchmark and evaluate the performance of various methods in deconvolution often necessitate the availability of sufficient ground truth data for comparison. Nevertheless, obtaining real bulk proportions experimentally can be challenging and constantly. Moreover, existing ground truth data tends to be limited in size and primarily centred around non-solid tissues. To facilitate method comparisons and understand their limitations, the generation of pseudo-bulk data proved to be a valuable approach. It should be noted that creating realistic simulations in deconvolution studies is essential for applying them confidently afterwards to real data. A common method to simulate bulk profiles involves utilising publicly available single-cell data and generating pseudo-bulk by averaging the expression of single cells. Simulation strategies employed in previous studies range from random sampling to more sophisticated approaches (Avila Cobos *et al*., 2020b; Dietrich *et al*., 2022; Hu and Chikina, 2023). Nevertheless, it is non-trivial to understand how the different parameters used in the simulations can affect the deconvolution results. In response, we have designed and compared various simulation approaches, introducing standardised parameters for simulating bulk expression profiles that can be used for the benchmarking of existing, and development of new methods.

#### Random sampling

First, we implemented a classic random sampling pseudo-bulk approach (simulation #1). In this simulation approach the parameters explored are: (a) the number of individual cells sampled for each pseudo-bulk (n) and the total number of generated pseudo-bulk samples (pool size). We systematically varied the pool size across three categories: 100 cells, 10,000 cells and 100,000 cells. At the same time, we explored the impact of the number of samples, conducting simulations with 100, 500 and 1,00 samples.

To explore the impact of the above parameters we performed random sampling with replacement using a single cell 10X dataset from mouse brain (Hrvatin *et al*., 2018a) composed of ∼48,266 cells and 28 samples (Supplementary Figure 1a-c). We observed that in this simulation, the number of cells sampled per sample affects the overall similarity of the distributions more than the number of samples (s). Moreover, we show that in the random sampling scenario the pseudo-bulks created are highly correlated and become almost identical as the number of sampled cells approaches the number of cells in the single-cell (Supplementary Figure 1d-e). However, it is crucial to note that real-world scenarios deviate from such uniformity. Bulk samples within a dataset might have extremely different proportions from sample to sample. In addition, various samples may lack certain cell types, introducing a complexity that is not fully captured in the random sampling simulations. Moreover, when the number of cells sampled from a single-cell dataset is significantly smaller than the total cell count (e.g 100 versus 22,000 cells), the simulation may only incorporate a sparse representation of each cell type. Moreover when we sample such a small percentage of cells per cell type, we might not capture rare cell types in our pseudo-bulk. It is essential to recognise the balance in random sampling - too much can lead to highly similar samples, while too little although it can yield diverse samples, they might fail to capture cell types effectively in terms of their signatures. It is crucial also to keep in mind when performing simulations that, in real bulk, samples typically comprise thousands to millions of cells.

#### Designing and comparing simulation methods

In our realistic simulations, we address limitations by introducing controlled noise. This noise originates from variances in proportions or the absence of cell types in pseudo-bulks. Simulation #2 involves random sampling using a uniform distribution (1-99) to select proportions, summed to 1. All cell types are sampled with replacement, resulting in a fixed range with large variance in proportions. To explore variability further, we introduce propVar in simulation #3, allowing us to adjust the uniform distribution range, producing arrays with more homogeneous to variable proportions. This parameter enables us to investigate the impact of proportion variability on deconvolution results. Additionally, we examine deconvolution methods in the presence of missing cell types in the bulk for simulation #3. A bimodal distribution with equal probability assigns cell types to mode 1 or 2. Mode 1 has a distribution with mean=0 and small standard deviation(sd)=0.0001, while mode 2 has mean=1 with variable standard deviation. This scenario mimics real-world situations where a tissue signature is known, but the presence of all cell types in the bulk is uncertain (Figure 2a). We finally compare the above simulation methods with random sampling as well as simulations proposed in other studies (Chu *et al*., 2022; Hu and Chikina, 2023).

**Figure 2.**
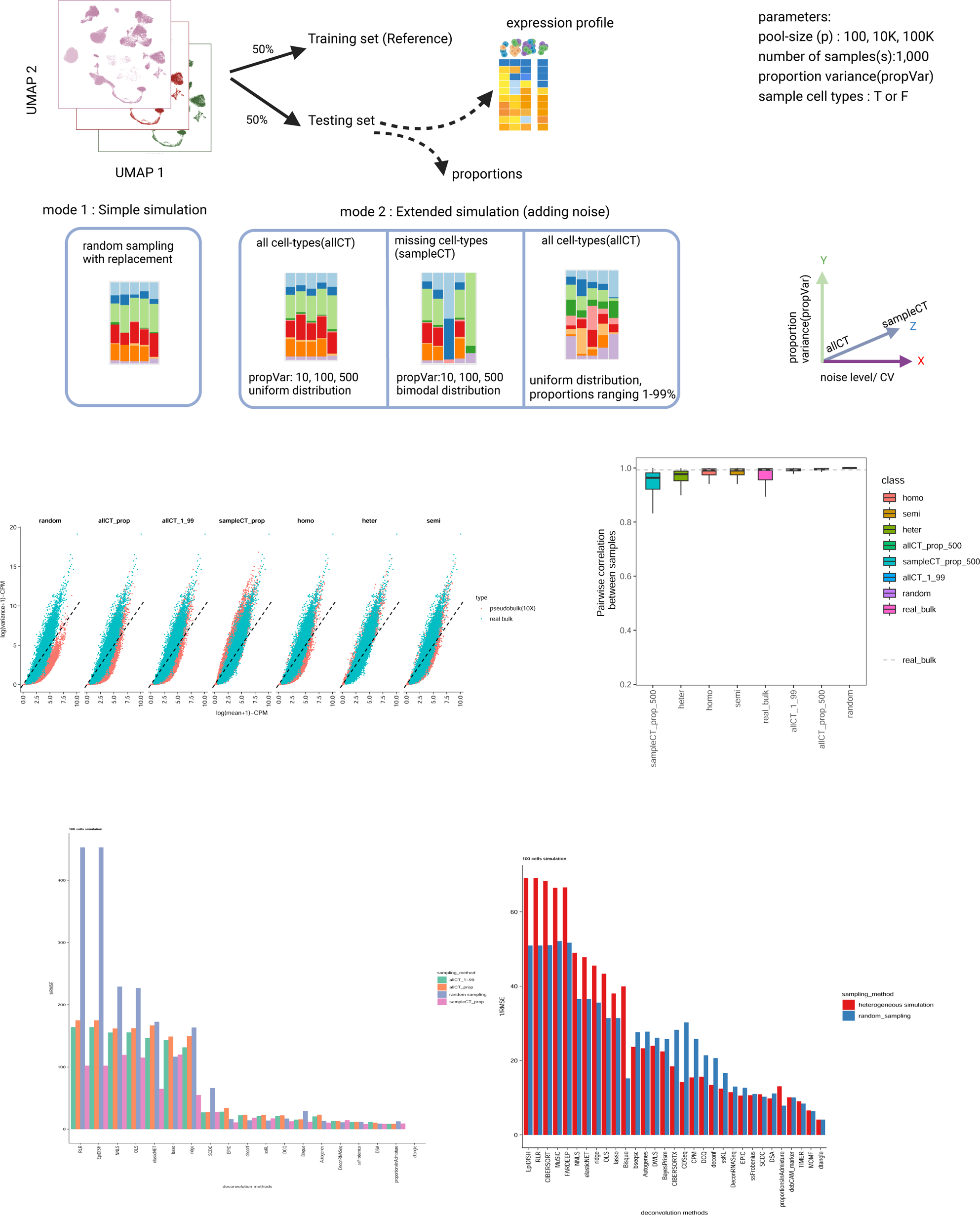
Evaluating pseudo-bulk simulations for benchmarking and new method development. a. Simulation workflow: each single-cell matrix is split into half to generate a training set (reference) and a testing set (pseudo-bulk matrix). 4 different sampling techniques were used to generate pseudo-bulks and ground truth proportions. We set 4 different parameters for the simulation: the number of cells to be sampled from the single-cell (pool size (p)), the number of samples in each pseudo-bulk matrix (s)-the default is 1,000 samples, the variance of the proportions (pVar) for the cell-types and finally a logical vector to decide if we use all the cell-type or not (sampleCT, True or False). b. Scatter plot of mean and variance of expression data from 7 different simulation scenarios and matched real bulk data. All the simulated pseudo-bulk data derive from the same mouse 10X single-cell dataset and the real bulk RNA-seq derives from the same study as well (Hrvatin2018). c. boxplot plot showing the pairwise correlation of gene expression for the samples of each simulation and for the real bulk. d. root mean square values (RMSE) of the 4 different simulation scenarios across the tested deconvolution method that were feasible to obtain results(10K cells simulation). e. Root mean square values (RMSE) of random sampling and ‘heterogeneous’ simulation across the tested deconvolution method that were feasible to obtain results(100 cells simulation).

#### Results and Conclusions

To understand the differences across simulations we compare pseudo-bulk gene expression with 10 real bulk mouse samples derived from the same study (Hrvatin *et al*., 2018b). One way to compare different bulk profiles is to calculate the mean against the variance of gene expression in logarithmic scale or the coefficient of variation (CV). Real bulk and generated pseudo-bulk samples exhibit a negative binomial distribution, as expected, for expression data (Robinson and Smyth, 2007) however, random sampling pseudo-bulk deviates the most from real bulk, with less overdispersion observed (Figure 2b). Moreover, when we compare the Coefficient of Variation (CV) in the simulated bulk against the CV in the real bulk, we observe that random, allCT_1_99 and allCT_propvar_500 are a lot less variant, sampleCT_prop_500, homogeneous and semi demonstrate higher CV, while heterogeneous simulation seemed to be closer to real bulk expression CV (Supplementary Figure 2a). At the same time we compared the methods based on the pairwise similarity of gene expression similar to Chikina et al (Supplementary Figure 2b), showing that heterogeneous and random simulation yield show lower gene correlation. We also compared the similarity of gene expression between samples in each pseudo-bulk strategy. A random sampling of 10K cells, and sampling all the cell types with variable proportions with replacement generates extremely similar pseudo-bulks while the other strategies produce samples with lower correlation, meaning that the simulated datasets using the latter approaches are more diverse and are closer to the bulk (Figure 2c). Our results suggest that random sampling from single cells fails to generate pseudo-bulks that mimic real bulk expression and therefore should be avoided when developing new methods, as it oversimplifies the deconvolution problem. This has also been discussed before and aligns with results from previous studies. On the other hand, sampling cell types creates pseudo-bulks that can better mimic real bulk expression variability and can challenge deconvolution methods to deal with greater noise in the bulk. Using all the cell types from single-cell with a medium variance of proportions (propVar=500, allCT_prop) provides a good starting point for evaluation and method development; while sampling cell-types (sampleCT_prop), semi and heterogeneous simulation represents the noisiest scenarios that can challenge deconvolution methods as demonstrated by the performance of deconvolution methods(RMSE values, Figure 2d). Many studies have talked about decrease in performance of methods when cell types are missing from the reference, here we show that this decrease is also happening when there are missing cell types in the bulk (sampleCT approach). Finally, we compared deconvolution performance in a mini-benchmark test between the random sampling and the heterogeneous method. We simulated 100 cells per sample for both methods and results demonstrate better performance of the heterogeneous method compared to the random sampling. This could result from the high variability depended on the small pool size (Figure 2e). More extensive benchmarks across the different simulations would be needed in the future. Overall, we conclude that pseudo-bulk simulation parameters affect deconvolution severely and developed methods should be tested for performance in a range of scenarios before applying deconvolution to real data.

### 3.3 Effect of pre-processing methods in deconvolution results

Previous studies have reported that pre-processing of input data in deconvolution impacts the deconvolution performance (Avila Cobos *et al*., 2020a; Jew *et al*., 2020), but it is yet unclear which methods should be used for each deconvolution method selected. Standard preprocessing of bulk and single-cell data includes the transformation and normalisation of the expression matrices. Transformation methods account for the mean-variance dependencies as well as the extreme count values present in a dataset. This is a crucial pre-processing step for most downstream analyses of both single-cell and bulk RNA-seq data. For instance, both the identification of DEGs and PCA analysis requires the transformation of raw count values. The same principle applies to single-cell data for performing dimensionality reduction and identification of marker genes. Gene expression normalisation is also an essential step for the above analyses. Normalisation methods correct gene expression matrices for factors that affect the number of reads that will be mapped to a gene (gene length, GC content and sequencing depth) (Evans, Hardin and Stoebel, 2017). In general, the goal of normalisation is to enable the comparison of gene counts within a sample and across samples. As mentioned before, deconvolution requires reference data that provides cell-type specific information in the form of single-cells, summed single-cells or marker genes derived from single-cell and bulk RNA-seq data to be deconvolved. Here we test how correcting data with transformation and normalisation techniques affect the deconvolution process. We implemented 2 transformation methods and 11 different normalisation methods in our pipeline (Table S2). Previous studies have applied transformation of the data first and then normalisation. Here we show that normalising the matrices first and then applying transformation yields better deconvolution results, on average in 8 brain tissues (Figure 3a-b, Supplementary Figure 3). We then examine the effect of different normalisation and transformation methods. Results from deconvolution in brain tissue show that the majority of methods output better results when the input data are not transformed. However, bseq-sc and BisqueRNA methods benefit from the logarithmic transformation of the input matrices (Figure 3c). Moreover, in many cases, sqrt-transformation seems to perform worse than no-transformation with the exception of the penalised linear regression methods lasso, elastic net and ridge where it performs the same as no transformation of the matrices. Lastly, we tested 11 different normalisation methods across 3 different brain datasets that have been produced from different single-cell technologies (10X and SmartSeq2). In all three datasets, row normalisation seems to perform worse across all the deconvolution methods (lower Pearson Correlation values), while the other normalisation methods appear to perform differently depending on the dataset. The majority of normalisation methods yield good results for the first dataset (Darmani2017). On the other hand, for the other two 10X datasets, none, global min-max, column-z score and mean normalisation methods appear to yield better results (Figure 3d-f).

**Figure 3.**
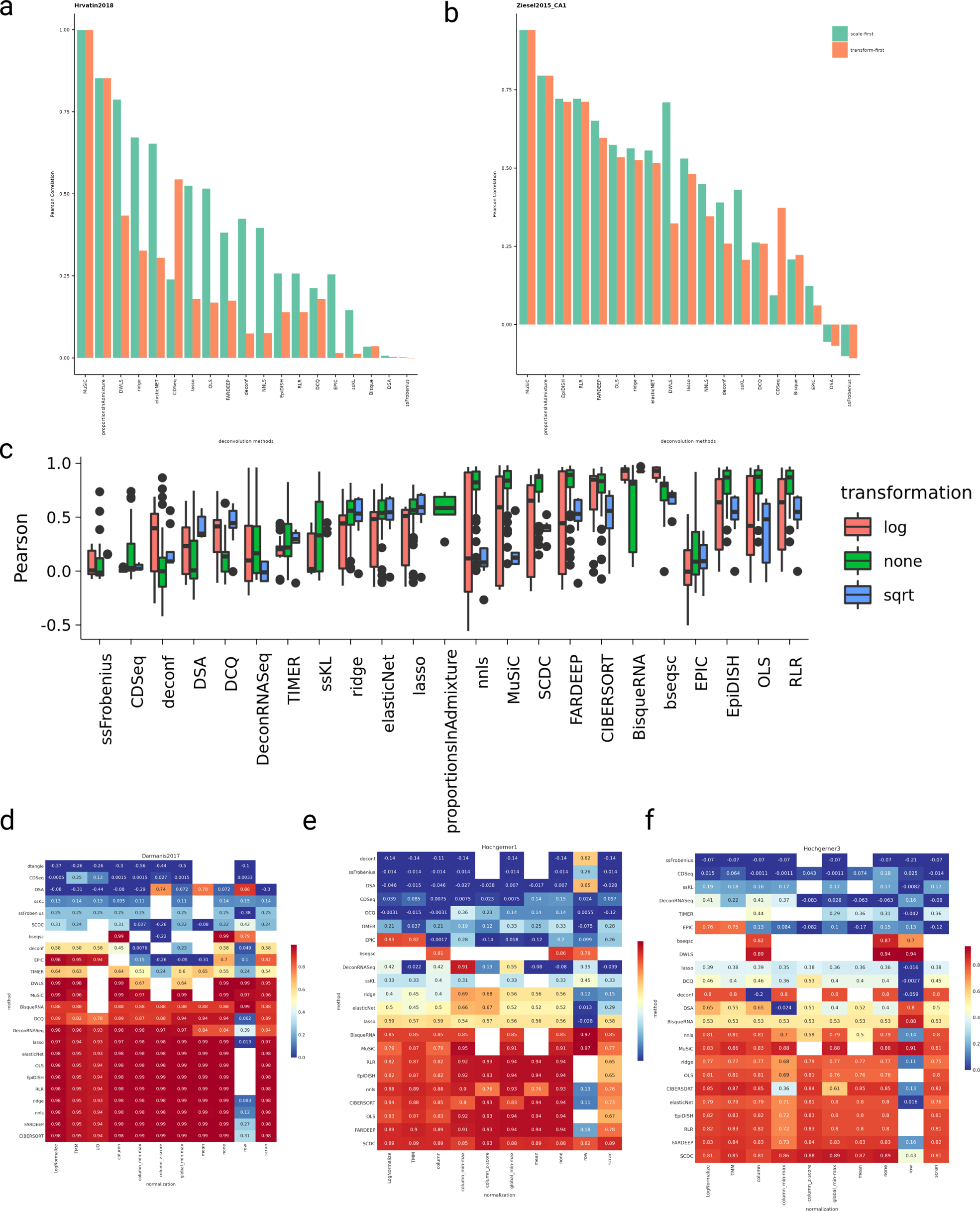
Pre-processing steps affect downstream decomposition of samples. Barplot showing Pearson Correlation values of a self-reference deconvolution task in which the effect of scaling first or transforming first the input matrices is tested a. Hrvatin 2018-brain tissue. b. Ziesel2015-brain tissue c. Boxplots showing Pearson correlation values estimated from the deconvolution of simulated data (from brain tissue) across 2 different transformation methods (log and sqrt) as well as no transformation (linear scale). Each data point in the boxplot represents a combination of a transformation and a normalisation method utilised in the deconvolution task (e.g log+TMM, log+column and similar for all the possible combinations). d-f. Results from deconvolution of 3 different simulated data from brain tissues (Darmanis2017, Hochgerner1, Hochgerner3) across 11 normalisation methods and 24 tested deconvolution methods.

### 3.4 Batch effects between (pseudo) bulk and single-cell data affect deconvolution

The first methods developed for deconvolution were utilising flow cytometry data from sorted cells, marker gene signatures or no references at all (unsupervised methods) to deconstruct bulk RNA-seq and microarray samples (Abbas *et al*., 2009; Gaujoux and Seoighe, 2013; Chen *et al*., 2018). Nowadays, with the advancement of single-cell protocols and the extensive availability of single-cell data from different species, a great opportunity has appeared to utilise them to facilitate deconvolution.

#### Bulk versus single-cell RNA-seq

Newly developed deconvolution methods are making use of single-cell data to deconvolve bulk profiles by selecting appropriate features (genes) such as MuSiC (Wang *et al*., 2019b), DWLS (Tsoucas *et al*., 2019)) and many others. Yet, the large differences between single-cell and bulk quantification methods of expression data can impact the outcome of deconvolution. Here, we compared the expression of each gene between single-cell data and real bulk data from the same tissue. We observed significant expression differences between summed single-cell data (pseudo-bulk) and real bulk data originating from the same tissue, here pancreas (Fadista *et al*., 2014; Baron, Veres, Samuel L. Wolock, *et al*., 2016) (Supplementary Figure 4a, Supplementary Figure 4b). In contrast, when comparing real bulk mixtures with each other, or pseudo-bulks with each other we observe very little differences (Supplementary Figure 4c-d). These discrepancies, observed between bulk and single-cell, are most likely due to the batch effects deriving from the different nature of the two sequencing techniques and dissimilar quantification methods. Principal Component Analysis (PCA) of the two data types, showed that PC1 could explain 74.5% of the variance between the bulk and the single-cell (Supplementary Figure 5a-c).

#### Self-reference versus Cross-reference deconvolution tasks

Next, we generated pseudo-bulk and single-cell references by randomly splitting the same dataset in half and then performing deconvolution afterwards. This task is a very simple case of deconvolution since both the bulk and the reference contain the same gene expression information. We name this ‘self-reference deconvolution’. While self-reference gives the opportunity to study deconvolution using simulations, it oversimplifies the problem. For this reason, we introduced cross-reference deconvolution tasks to more systematically study the impact of batch effects in cell-type deconvolution. For this task, we split the same pancreatic single-cell dataset in half and we selected cells from two individuals to generate the reference while the other two biological samples were used to build the bulk profiles. In this way, we artificially introduced batch effects in cellular decomposition. Deconvolution results from one “bulk” and one “single-cell” algorithm (FARDEEP and DWLS) show a significant drop in deconvolution performance in the cross-sample task compared to the equivalent self-reference task (Figure 4a, Supplementary Figure 6a). These results suggest that cell-type deconvolution is severely affected by batch effects caused by expression differences across samples. Furthermore, since the deconvolution of real bulk samples typically entails expression data from completely different studies and the use of different technologies, we next studied how cross-study and cross-technology tasks can impact the performance of deconvolution algorithms. For the cross-study deconvolution, we selected two human pancreatic single-cell datasets (Baron, Veres, Samuel L. Wolock, *et al*., 2016; Segerstolpe *et al*., 2016) (Table S1) which were generated from two different laboratories. We observed that deconvolution across studies can impact deconvolution performance severely when compared to self-reference tasks (Figure 4b, Supplementary Figure 6b). Next, we compared the performance of deconvolution when datasets from different technologies are involved. For this task, we focused on 3 independent dataset pairs from the placenta, brain and lung tissue (Supplementary Figure 7, Supplementary Figure 8).

**Figure 4.**
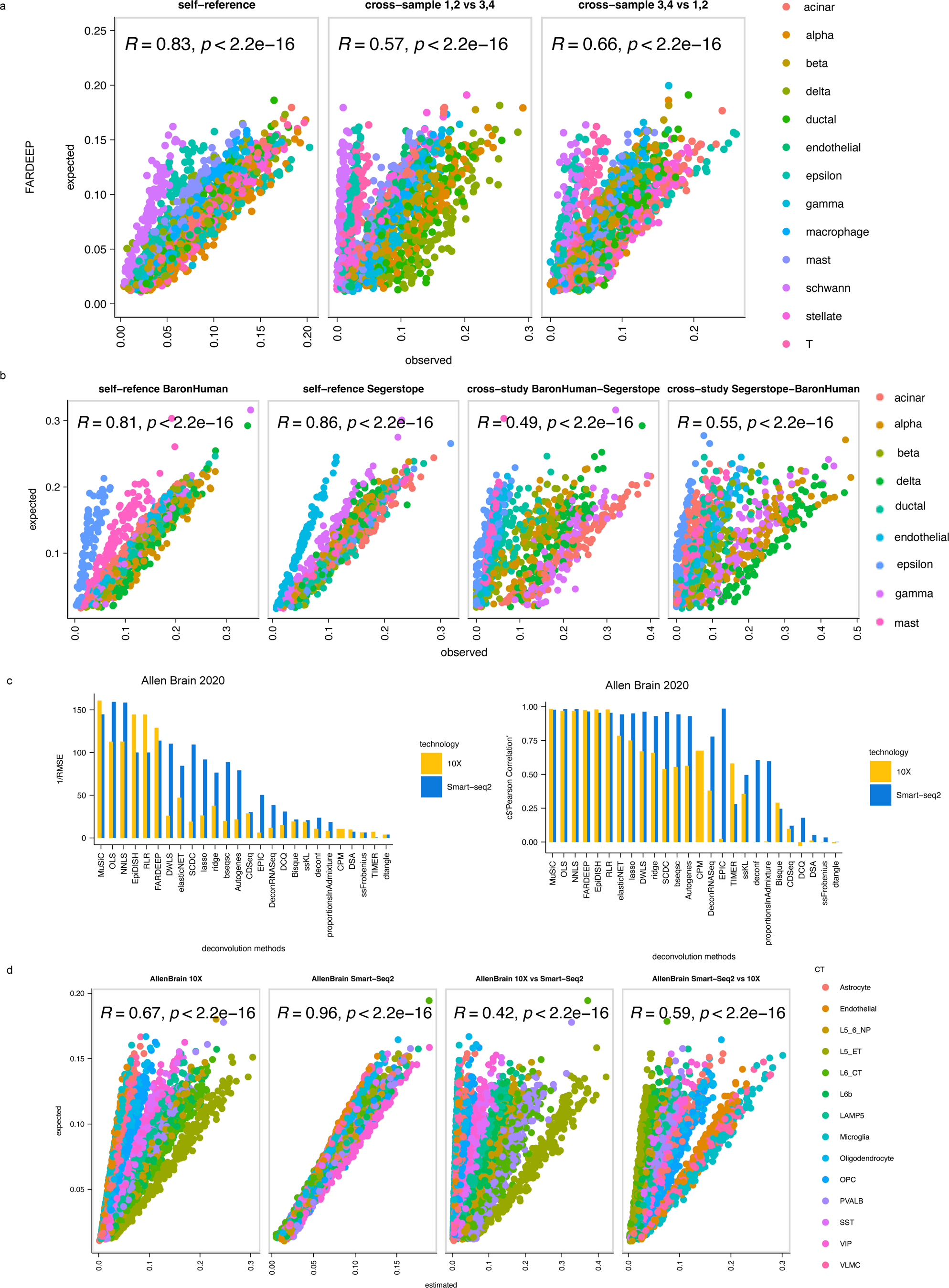
Impact of batch effects in deconvolution. a. scatter plots showing results from self-reference and 2 examples of cross-sample deconvolution from the method FARDEEP utilising a single-cell from human pancreas (13 cell-types). In the self-reference task all 4 samples are used while cross-sample tasks use two samples as reference and the other two represent the pseudo-bulk at each example. Pearson correlation (R) and p-value are reported for each task. b.scatter plots showing the results from two self-reference tasks from two pancreatic studies (9 cell-types) as well as two cross-study deconvolution tasks. c. Bar plots showing 1/RMSE and Pearson correlation values from the 26 feasible deconvolution runs from two human brain datasets that come from the same study but different technologies (10X and Smart-seq2). d. Scatter plots demonstrating results from the DWLS method from the two self-reference tasks (10X and Smart-seq2 of human brain tissues, 14 cell-types) and two cross-technology tasks. Pearson correlation (R) and p-values have been calculated for each task and shown on the plots.

Each dataset pair was generated from the same laboratory but using different single-cell technologies (10X and Smart-Seq2). Given that 10X and Smart-Seq2 are the most common single-cell protocols available, this test will help reveal which references are suitable in deconvolution. Results show that self-reference deconvolution with 10X sc-RNASeq data yields better results overall (higher Pearson correlation, lower RMSE values Figure 4c, Supplementary Figure 6c-d) across most deconvolution methods compared to Smart-seq2 self-reference. Moreover, results from the top method in this task (DWLS) demonstrate that cross-technology deconvolution results in less accurate predictions, with very low Pearson Correlation values observed (Figure 4d). All the above highlight the importance of the reference selection and the barrier that batch effects oppose to deconvolution.

### 3.5 Evaluation and consensus-based approach for deconvolution in real biological samples

#### Deconvolution with flow cytometry as ground truth

In contrast to the simulation tasks performed above, here we decompose real bulk profiles from 9 Peripheral Blood Mononuclear Cells (PBMC) (Finotello *et al*., 2019) samples, utilising our deconvolution pipeline to assess the results and generate an honest consensus of predictions of cell-type proportions. Apart from the publicly available RNA-seq dataset from the above study, there is also flow cytometry data available for the 9 individuals which can be used as ground truth in our deconvolution pipeline. The researchers have measured the proportions of 8 immune cell types using specific markers for each cell population (CD4+ T cells, CD8+ T cells, T-regulatory cells, B cells, NK cells, myeloid dendritic cells, monocytes and neutrophils (Supplementary Figure 9a,d). For this deconvolution task, we selected a recent single-cell PBMC dataset (Wilk *et al*., 2020) to use as a reference. We re-analysed the single-cell data and matched the cell types between the single-cell and the ground truth data from flow cytometry so that our reference has the same cell types that had been measured in the bulk (Supplementary Figure 10a-e). We also summarised the proportions of CD4+positive and T-regulatory cells measured by flow cytometry since our single-cell did not contain T-regulatory cells (Supplementary Figure 9b-c, Supplementary Figure 10e). Before we apply deconvolution on the 9 real bulk samples we first performed a self-reference deconvolution to evaluate how the reference performs in the pseudo-bulk deconvolution task. Nine methods yield good results with a Pearson Correlation of more than 0.86 (Supplementary Figure 10f). Next, we applied the all 31 deconvolution methods included in the pipeline on the real bulk PMN/PBMC data and obtained the cell-type predictions. The methods were evaluated across 9 evaluation metrics. Pearson Correlation, Spearman Correlation, RMSE and weighted RMSE values are the main metrics we focused on in this benchmark task (Figure 5). Results from the additional metrics implemented in the pipeline are shown in Supplementary (Supplementary Figure 11). DWLS, OLS and bseq-sc showed the highest Pearson Correlation values while DWLS and bseq-sc also maintained good RMSE and Weighted RMSE values (Figure 5). Looking closer at the DWLS cell-type predictions we observe differences across samples (Supplementary Figure 12a-b). Moreover, we examined the deconvolution predictions per cell type and noticed that CD4+ cells were consistently overestimated while CD8+ cells were underestimated in all 9 donors by the DWLS deconvolution method. The same pattern can be validated with the bseq-sc method which was ranked second in this task based on Pearson Correlation values (Supplementary Figure 12). It has been reported before (Aliee and Theis, 2021) that it is difficult to deconvolve closely related cell types due to collinearity, a problem which some single-cell methods attempt to solve with sophisticated feature selection. Since the large benchmarking across all the deconvolution methods has been performed using all the genes as initial input (except for the marker specific methods (e.g DSA, deCAMmarker, dtangle) we explored how different signatures affect deconvolution results of marker-based and reference-based methods. For this analysis we compared deconvolution results using all the genes, different marker selection algorithms(MAST, wilcox test, t-test) and LM22 immune signature. Results show that many methods perform well when LM22 signature is used but their performance decreases when other sets of signatures are selected. Nevertheless methods such as OLS, EpiDISH, RLR, CIBERSORT and other seem to be more robust in the selection of different signatures (Supplementary Figure 13a-b)

**Figure 5.**
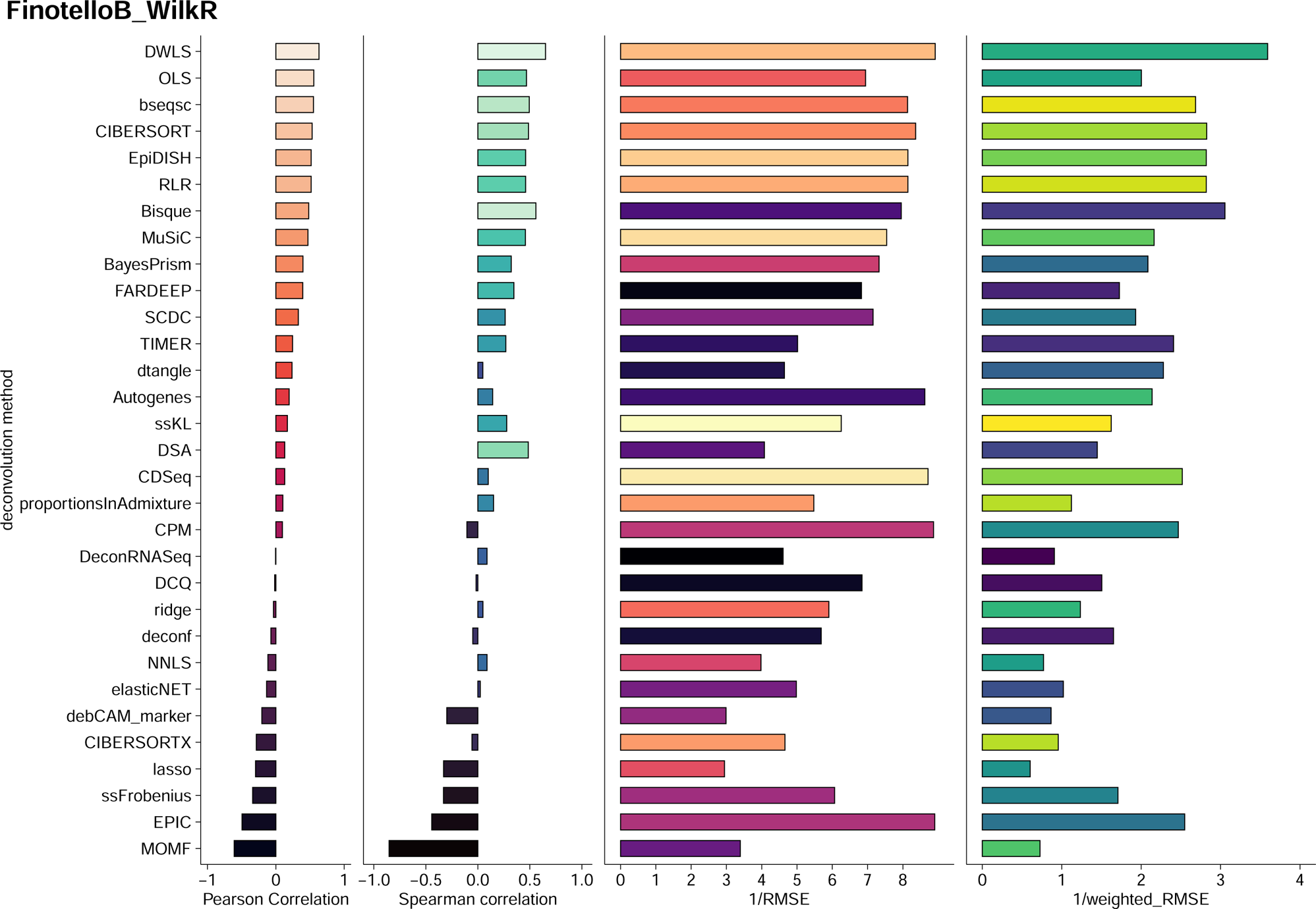
Deconvolution performance of real bulk RNA-seq samples. Results from 4 main metrics (Pearson Correlation, Spearman Correlation, RMSE and weighted RMSE for 31 deconvolution methods on real bulk PBMC data with ground truth using a single-cell PBMC dataset as reference (Wilk2020).

#### Deconvolution with tissue slides as ground truth

Since there is no single method that clearly ranks first in all scenarios and it is still unclear how different methods work in data from different tissues, we believe that a fair deconvolution consensus is a reasonable approach to take. Here, we suggest a consensus method based on 3 methods DWLS, FARDEEP and EpiDISH (selection criteria on Methods) that perform well in previous pseudo-bulk scenarios and compute proportions relatively fast and reliably (Supplementary Figure 10). Bseq-sc although it performs well, because of its iterative nature, it can be very slow (> 1 week) to compute proportions so it was excluded from the consensus calculations. Next we applied the consensus on GTEx stomach data (339 samples) for which we had access to tissue slides that could inform the validation of the deconvolution results. For this task we used as a reference the stomach subset of the Human Cell Landscapes sRNA-seq dataset containing 16 cell-types (Figure 6a-b). From the deconvolution results we can observe a clear difference between the samples on the left of the barplot and the right hand of the barplot. Samples on the right seem to contain mostly smooth muscle cells, fibroblasts and endothelial cells, whereas samples on the left are mainly peptic and goblet cells (Figure 6c). Next we explored if this difference in the composition can be observed in the tissue slides of the adjacent samples. We randomly selected a small number of tissue samples (5 from the samples from the left side and 5 from the right side). We observed that the slides that come from samples from the right side of the barplot, were in general more enriched in muscle cells compared to the samples on the left side which were more enriched in mucosa and submucosa tissue which aligns with the presence of goblet and peptic cells in the deconvolved mixtures (Figure 6e,f). We also quantified the mucosa, submucosa and muscle layers(μm2) for each stomach sample using QuPath and computed the proportions of each layer in each sample. Barplot showing the proportions computed for the 339 samples in the sample ordered the same way as in Figure 6c. Although we can observe that the samples on the right have a lot less mucosa and more muscle which aligns with the deconvolution results (absence of peptic and goblet cells on the samples on the right and presence of smooth muscle), the measurements from the tissue slides seem to be quite different from the deconvolution results overall. This could be because of a number of reasons: (a) annotating mucosa,submucosa and muscle layers is not straightforward and is dependent on the individual who performs the annotation. (b) deconvolution predictions are objected to overfitting (c) tissue slides might have different composition compared to the bulk sample that goes to sequencing due to sampling reasons. More detailed analysis on the comparison between composition in bulk samples and tissue slides should be made to conclude if tissue slides are a good ground truth data. To further explore the differences between deconvolution results and composition calculations from tissue slides we mapped each cell-type to the respective layer based on literature knowledge (see Methods) and we performed Pearson correlation analysis. We observe the highest positive correlation values between the Mucosa proportions from tissue slides and the summarised proportions of mucosa cells from deconvolution and similar Pearson correlation values between the Muscle layer and the summarised proportions from the cell types in the muscle layer. However the submucosa proportion from the tissue slide is not strongly correlated with any of the cell type proportions from deconvolution.

**Figure 6:**
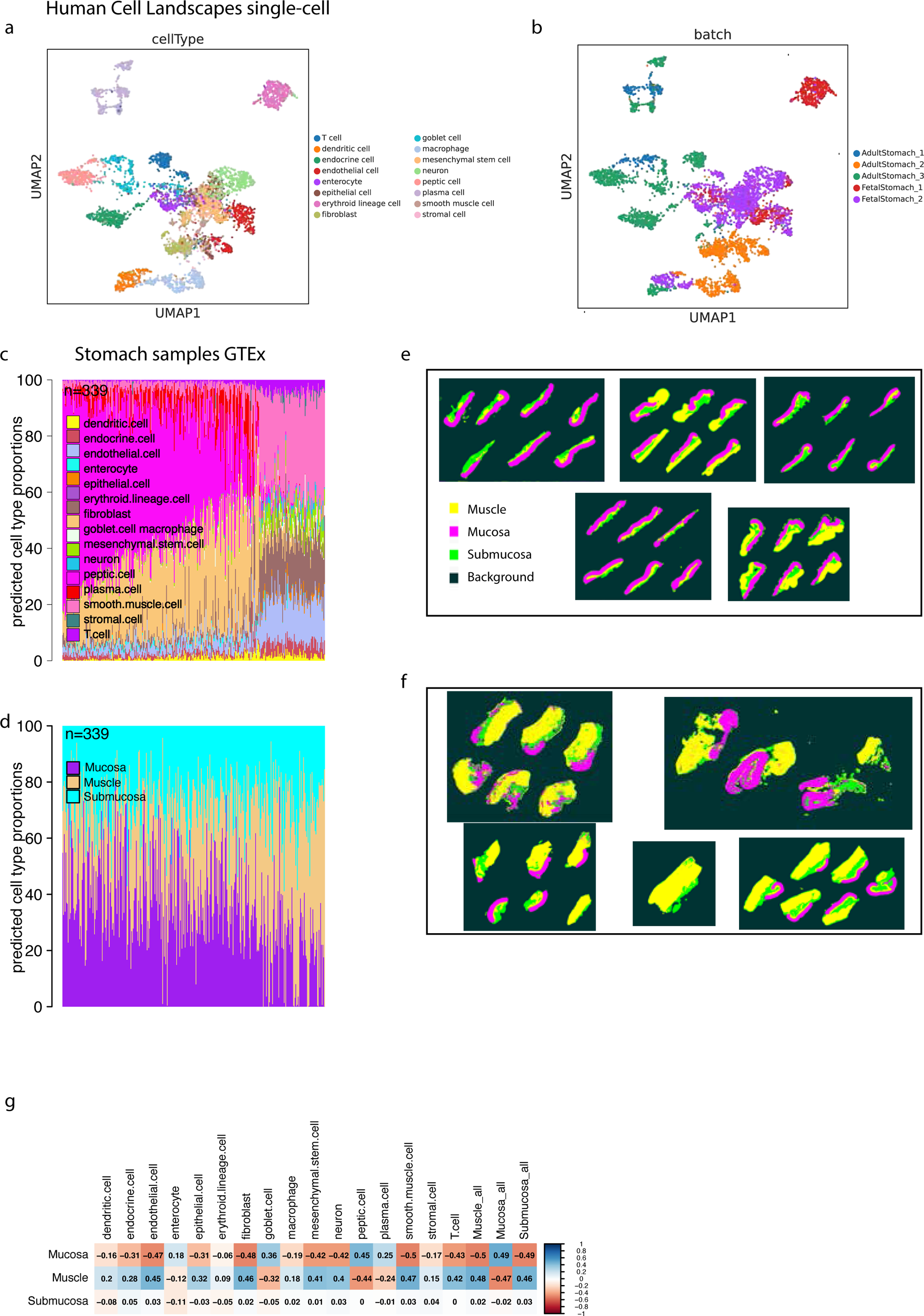
Application of consensus approach on stomach GTEx bulk expression data. a,b UMAP plots of the stomach subset of Human Cell Landscapes single-cell reference used for the stomach GTEx deconvolution showing the different annotated cell-types and the distribution of donors in the datasets. c. Bar plot showing the prediction of cell-type proportions from the consensus approach across 339 selected stomach samples that have matched tissue slides. d. Quantification of the different stomach layers (mucosa, submucosa, muscle) from the matched tissue slides using QuPath. Area μm2 is measured for each layer in each tissue slide. e. 5 randomly selected tissue slides from the samples on the left side of the barplot c. Submucosa is shown in green, mucosa in magenta and muscle compartment in green f. 5 randomly selected tissue slides from samples originating from the right side of barplot c g. correlation plot showing the Pearson Correlation values between the proportions measured from tissue slides and the predicted proportions from deconvolution of the bulk RNA-seq samples

#### Background prediction using experiments from sorted cells

Next, we designed an experiment in order to evaluate the background prediction of the consensus method in two datasets with purified cell types and/or mixtures with known cell types. Our data collection included mouse cerebral cortex and human lung datasets, for which we carefully selected relevant single-cell references tailored to the task (Supplementary Figure 14a,e). Initially, we applied the consensus deconvolution method to 17 samples obtained from the mouse cerebral cortex (Supplementary Figure 14b). Each sample in this task represented a single cell type, serving as the ground truth, and aimed to evaluate the method’s ability to accurately predict the presence of that specific cell type while quantifying background prediction. To quantify background prediction, we employed logarithmic loss, a widely used metric for evaluating classification model performance. In the context of deconvolution, this metric measures the accuracy of estimated cell type proportions within a mixture, taking into account uncertainties and penalising incorrect estimates arising from low-level gene expression that may lead to background predictions. Across the 17 samples, we observed consistently low log-loss values, ranging from approximately 0.004 to 0.104 (Supplementary Figure 14c). Notably, among all samples, two endothelial flow cytometry samples and one myelinating oligodendrocyte sample exhibited the lowest log-loss values, signifying precise predictions of the purified cell type within each sample. Moving forward, we extended our analysis to encompass 26 samples from human fetal lung. Of these, the initial 16 samples were predominantly composed of endothelial cells, while the subsequent 20 samples contained non-endothelial cells. Our objective was to observe whether the consensus approach could effectively differentiate between these two sample categories. Notably, the endothelial samples predominantly exhibited high proportions of endothelial cells in the predictions, albeit with predictions of fibroblasts, lymphocytes, and myeloid cells. In contrast, the non-endothelial samples were primarily predicted to consist of fibroblasts, with minor proportions of endothelial cells detected. These low-level endothelial cell predictions may be attributed to background predictions generated by the consensus algorithm (Supplementary Figure 14d).

#### Deconvolution in the absence of ground truth

In situations where ground truth data is unavailable, our pipeline incorporates a separate feature to address this. We perform a pairwise comparison of the deconvolution results across all samples, enabling us to estimate Pearson Correlation coefficients. These coefficients serve as a quantitative measure of the agreement between different methods. The results of this comparison are visually presented in the form of a heatmap, where methods that yield similar results cluster together (Supplementary Figure 10g). This clustering provides valuable insights into which methods exhibit strong agreement, both in the presence and absence of ground truth data.

It is important to mention that while the clustering approach successfully identifies method agreement, it does not inherently imply that the clustered methods consistently perform well. Finally, to comprehensively assess the reliability and scalability of our pipeline, we conducted parallel evaluations of each module, encompassing pseudo-bulk generation, normalisation, and deconvolution methods. These assessments provide valuable insights into the computational efficiency of our pipeline, reporting essential metrics such as processing time, memory usage, mean load, and CPU time for each module(Supplementary Figure 15a-b).

## 4. Discussion

In this study we developed a deconvolution framework that allows the evaluation of 31 publicly available methods across different scenarios with both synthetic and real data. We first studied the effect of different pseudo-bulk techniques by defining different ways of pseudo-bulk sampling. Our results demonstrated that pseudo-bulks generated with different methods produce highly diverse results in deconvolution, which shows that the results of deconvolution are highly sensitive to the sampling of cells and the selection of proportions. Here, we suggest the use of pseudo-bulk generation methods that can challenge deconvolution methods and reflect real bulk data, when developing new, or benchmarking existing methods. Another important aspect of deconvolution, that we explore, is the normalisation and the transformation of the input matrices in deconvolution. Both bulk and single-cell matrices used in the deconvolution pipeline were tested across different normalisation and transformation methods. Results suggest that normalisation can be beneficial in deconvolution since it corrects for differences in the library size both in the single cell and the bulk RNA-seq data. On the other hand, logarithmic transformation of the input matrices can result in worse performance with the exception of two deconvolution methods (bisqueRNA and bseq-sc). Similar to the reports from previous studies, this suggests that keeping input data in linear scale aids in assessing the cell proportions accurately (Avila Cobos *et al*., 2020a; Jin and Liu, 2021).

Another key aspect of the bulk sample deconstruction is the selection of the reference matrix, which is used to extract the key features (genes) that will be used in deconvolution. Results from Smart-seq2 and 10X single-cell datasets that have been used in this study, show that 10X data is consistently performing better as a reference, this is likely because this technology captures more cells of a specific cell type and as a result, more accurate features are extracted from these data. It should also be noted that in tasks when the bulk and the reference data come from different sources (studies or technologies) - the accuracy score of the deconvolution drops significantly. This indicates that differences in the expression of genes, across datasets, affect deconvolution heavily. Methods that take into account and successfully minimise these effects (DWLS, MuSiC) seem to also perform better overall in cross-reference tasks. We apply deconvolution in real PBMC samples using a suitable matching single-cell reference and conclude that two methods, DWLS and bseqsc perform best. All the above steps that should be performed before deconvolution have been implemented as part of the pipeline with a number of parameters that can be defined by the user.

Notably, in the absence of ground truth data, our pipeline introduces a valuable feature for pairwise comparisons of deconvolution results across all samples, allowing the estimation of Pearson Correlation coefficients. This innovative approach offers a quantitative measure of method agreement, even without the presence of ground truth data, providing researchers with insights into method performance. It is imperative to understand that while this clustering technique effectively identifies method agreement, it doesn’t inherently imply consistent method performance. We also suggest a consensus method based on three robust techniques: DWLS, FARDEEP, and EpiDISH and test it across 3 real bulk datasets with available ground truth testing its accuracy and background prediction.

Notably, the CATD pipeline, developed using snakemake, enables the reproducibility of our results and can be additionally utilised by users to deconvolve new bulk samples by providing a bulk dataset and a single-cell dataset from the same tissue. When ground truth is provided too, the pipeline will provide evaluation metrics for each method. On the other hand when ground truth is absent the pipeline will perform pairwise comparison of the results and report a heatmap with methods to be selected as well as calculating proportions based on the consensus method.

Moreover, the advent of new high-quality single-cell atlases with standardised annotation from both healthy and disease conditions will provide more opportunities to explore the contribution of cell proportions in pathological conditions. Finally, the development of robust, accurate tools for feature selection that enable clear distinction between cell-types will pave the way for use of computational deconvolution perhaps even in clinics or in settings where scRNA-Seq is not feasible.

## Authors Contributions

Anna Vathrakokoili Pournara: Conceptualization, Data curation, Formal Analysis, Investigation, Methodology, Software, Validation, Visualisation, Writing-original draft.

Zhichao Miao: Conceptualization, Data Curation, Methodology, Software, Supervision, Writing-review & editing.

Ozgur Beker: Methodology, Formal Analysis, Software

Nadja Nolte: Data curation, Formal Analysis, Investigation, Methodology, Software, Validation, Visualisation

Alvis Brazma: Conceptualization, Supervision, Writing -review & editing

Irene Papatheodorou: Conceptualization, Project Administration, Supervision, Writing-review & editing

## Acknowledgements

The authors would like to thank the reviewers for their valuable feedback and suggestions for the manuscript. The authors would also like to thank Craig Russell for his feedback and help in reviewing and editing the manuscript.

## Funding

This work was supported by: the European Molecular Biology Laboratory (A.V.P, Z.M., O.B.,N.N, A.B, I.P.); the EMBL international PhD program (A.V.P.) and the OpenTargets (N.N,C.M, I.P,) [OTAR2067]

## Conflict of interest

The authors declare no competing interests

## List of Supplementary Tables

**Supplementary Table 1:** Information on single-cell RNA-seq datasets from brain, placenta, blood, pancreas and lung tissue

**Supplementary Table 2:** Normalisation methods implemented in the CATD pipeline

**Supplementary Table 3:** Deconvolution methods implemented in the benchmarking pipeline CATD

**Supplementary Table 4:** Summary of real bulk data deconvolution from 4 different tissues using single-cell references

## Notes

https://github.com/Papatheodorou-Group/CATD_snakemake

### Competing Interest Statement

The authors have declared no competing interest.

### Summary of Updates

Included two additional methods in the pipeline(BayesPrism and CIBERSORTx); Included additional simulation methods comparison.: Additional metrics were included in the benchmarking(weighted RMSE and Spearman Correlation). Introduction and Methods sections have been updated. Introduction section: submitted a correction for the use of enrichment-based methods.

