## Supplementary figures and images for "CATD: A reproducible pipeline for selecting cell-type deconvolution methods across tissues"

### Supplementary Figure 1

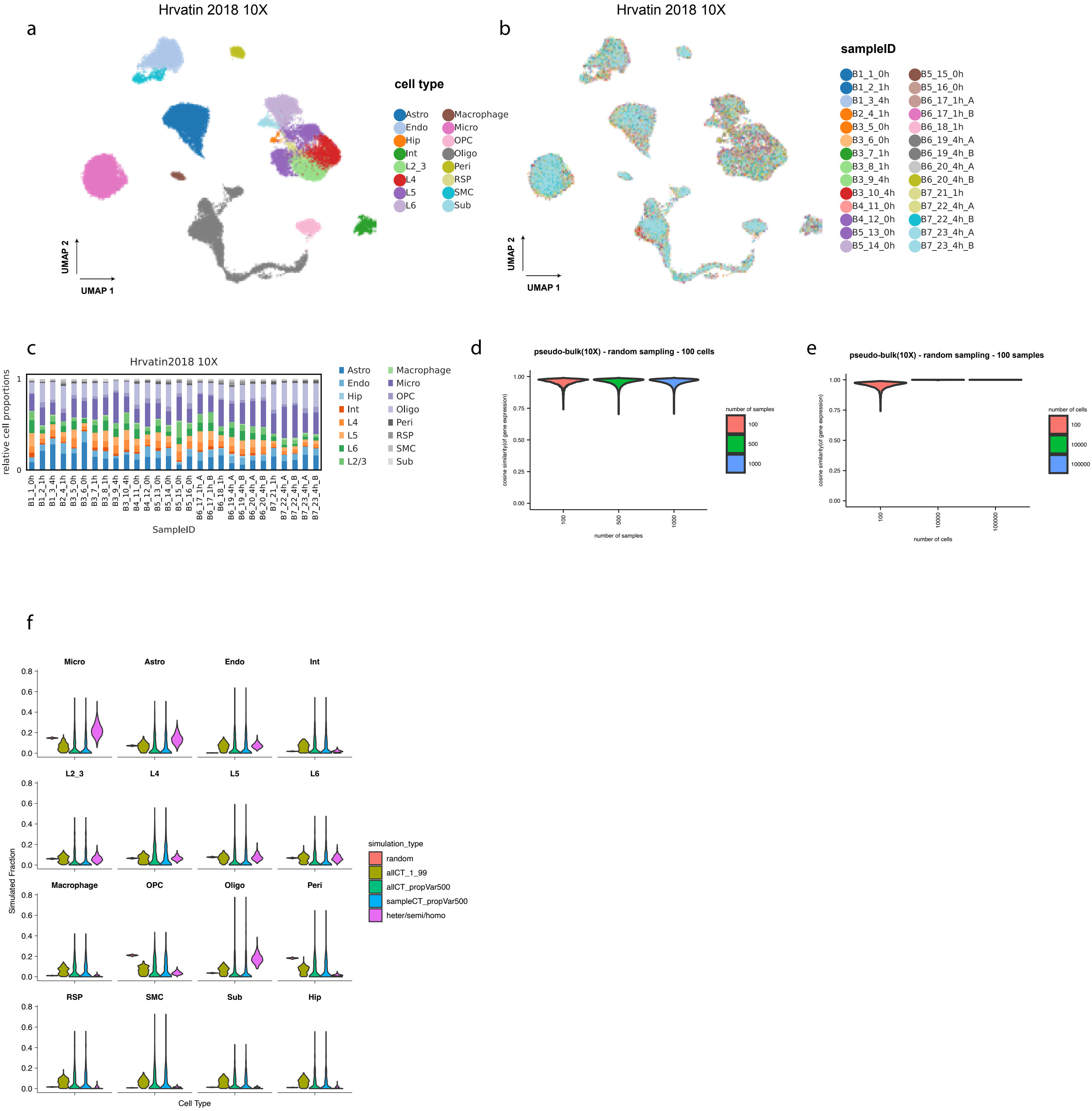

### Supplementary Figure 2

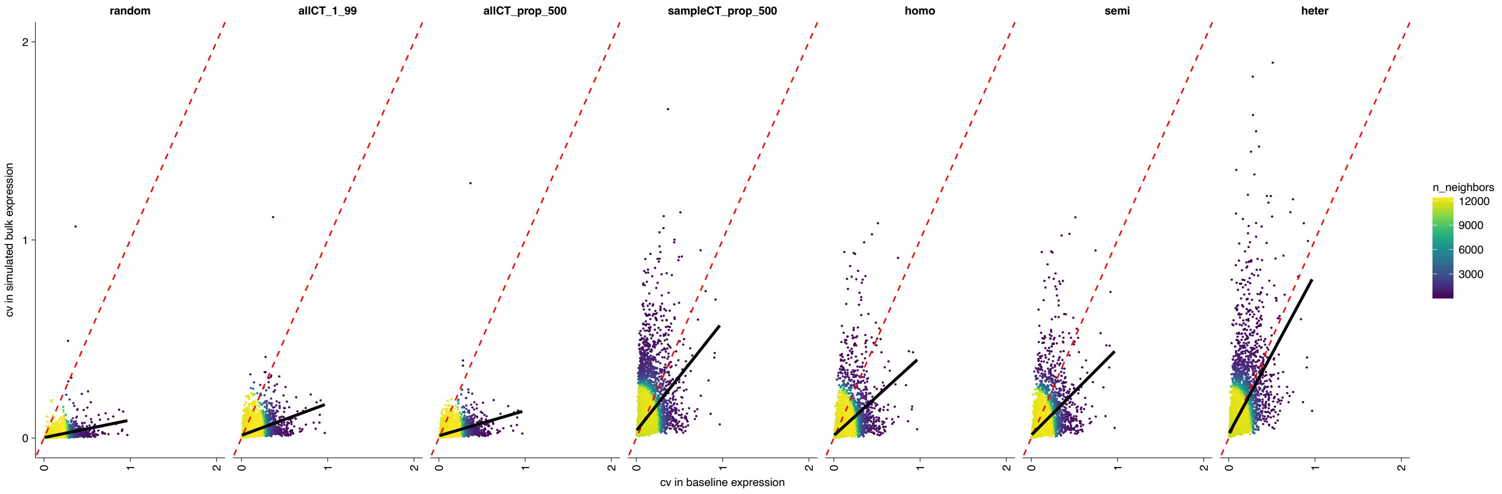

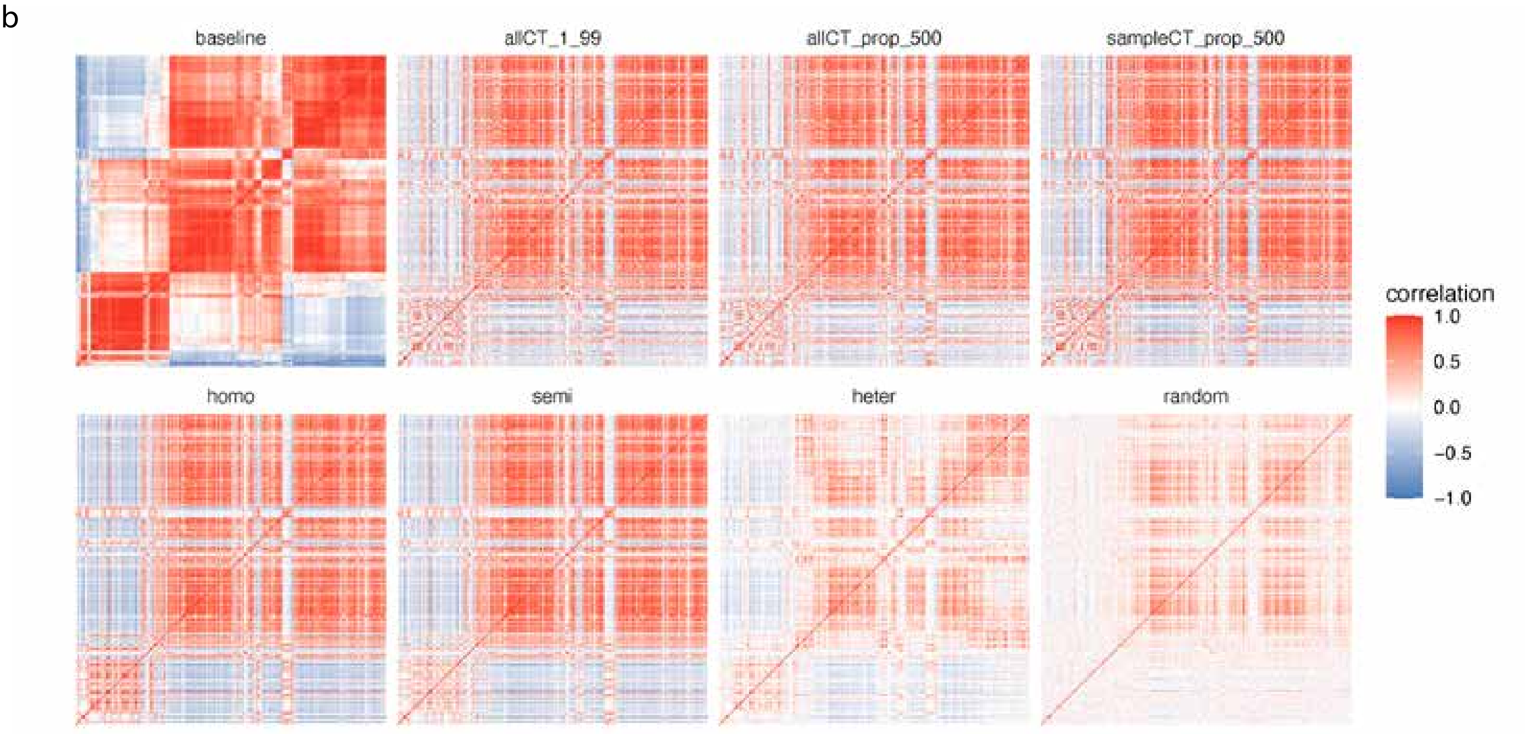

### Supplementary Figure 3

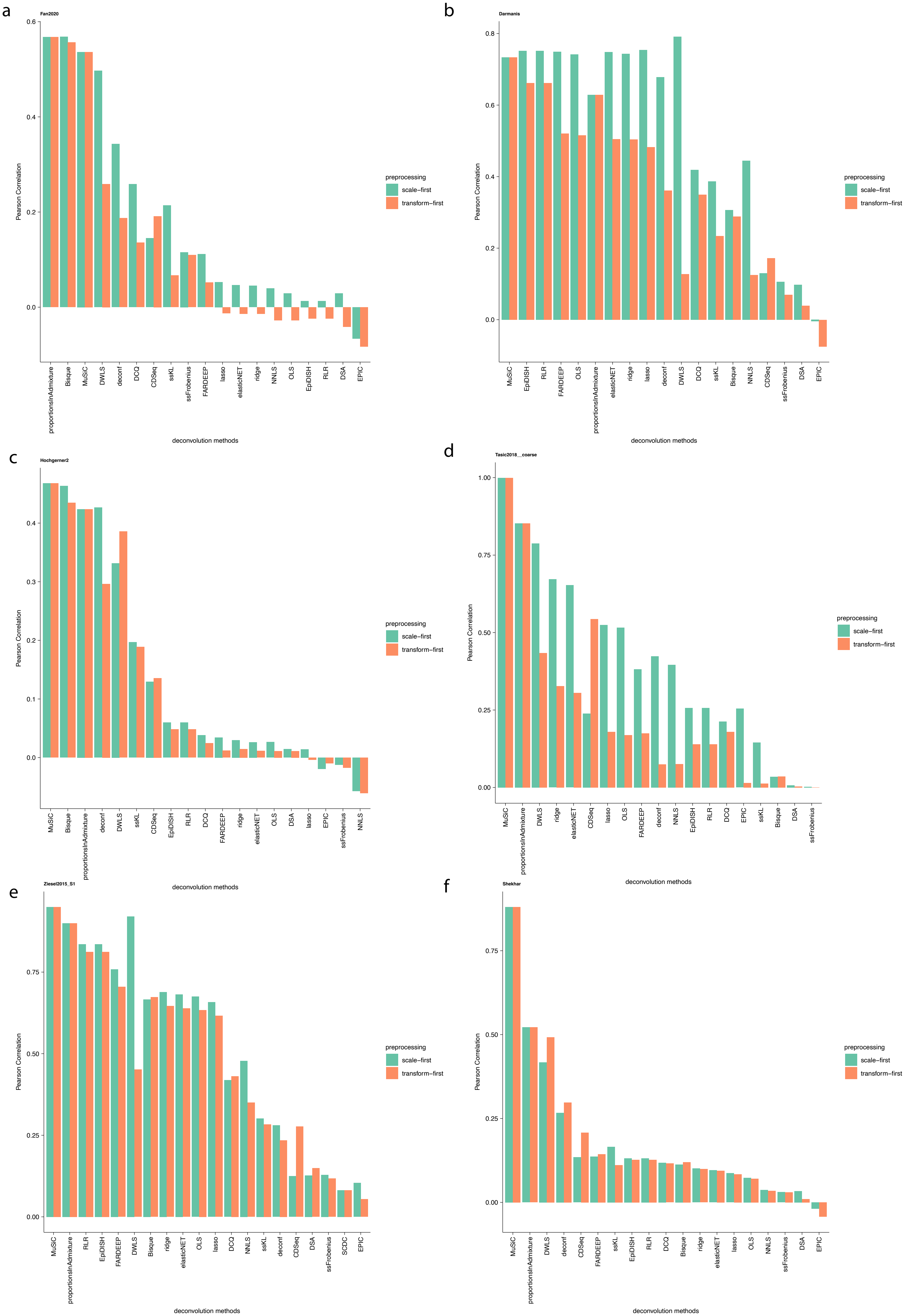

### Supplementary Figure 4

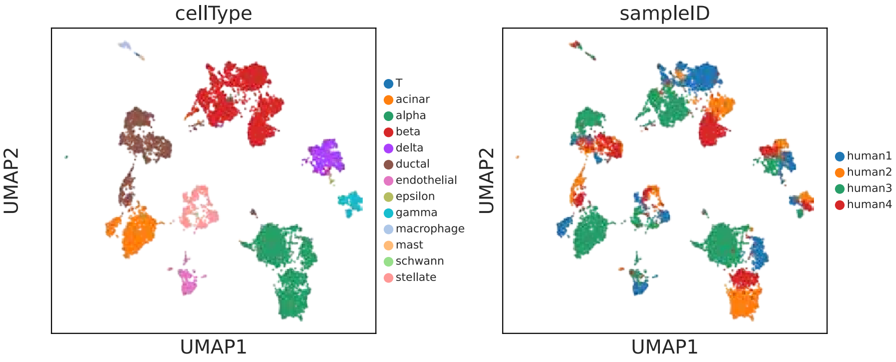

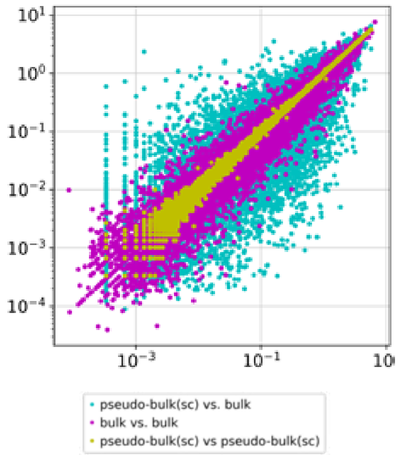

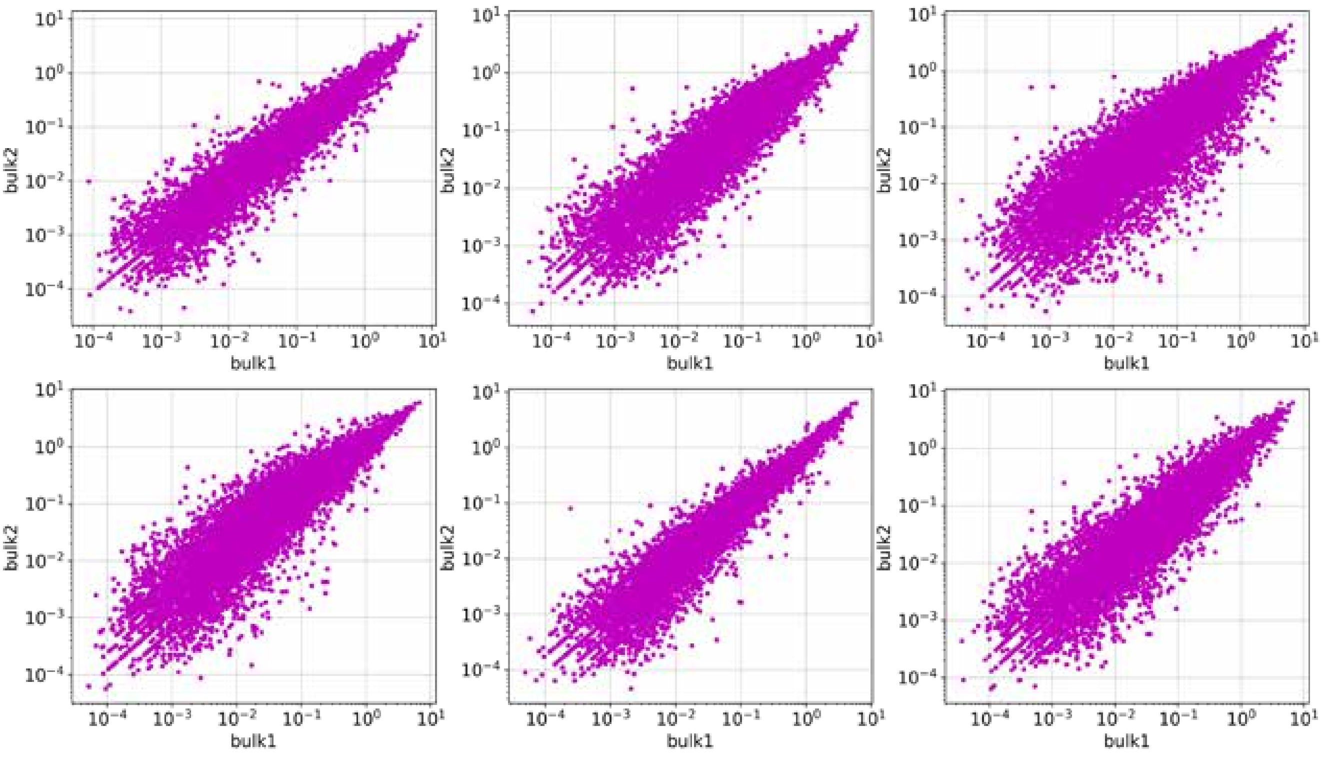

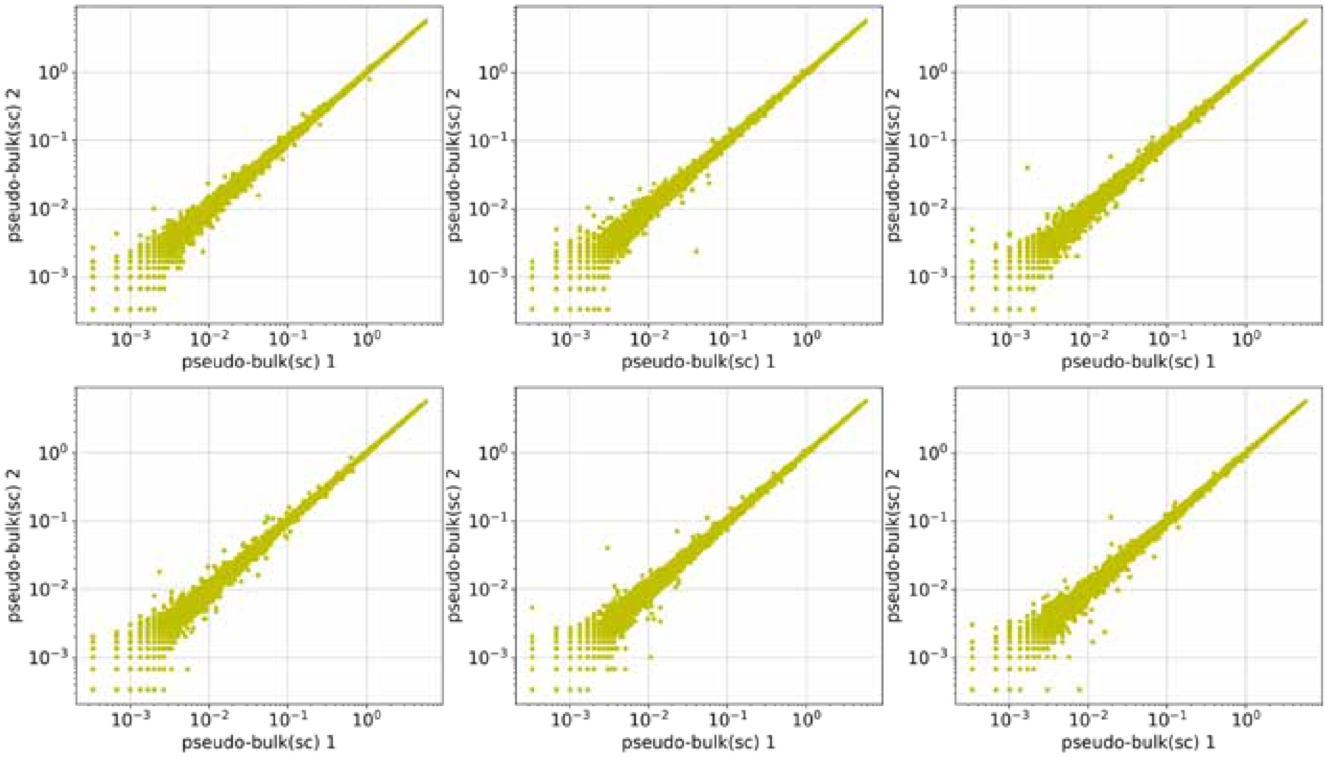

### Supplementary Figure 5

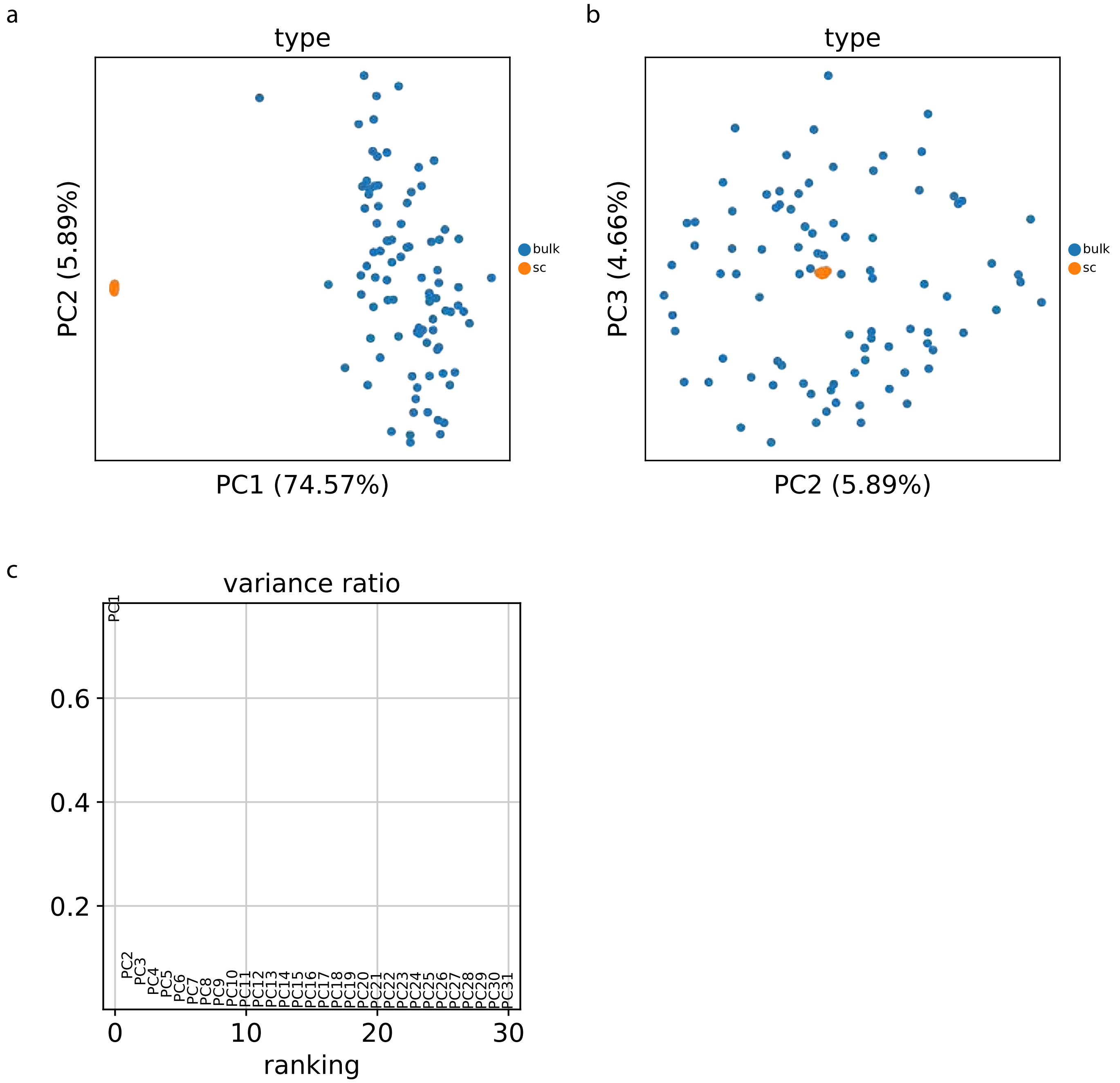

### Supplementary Figure 6

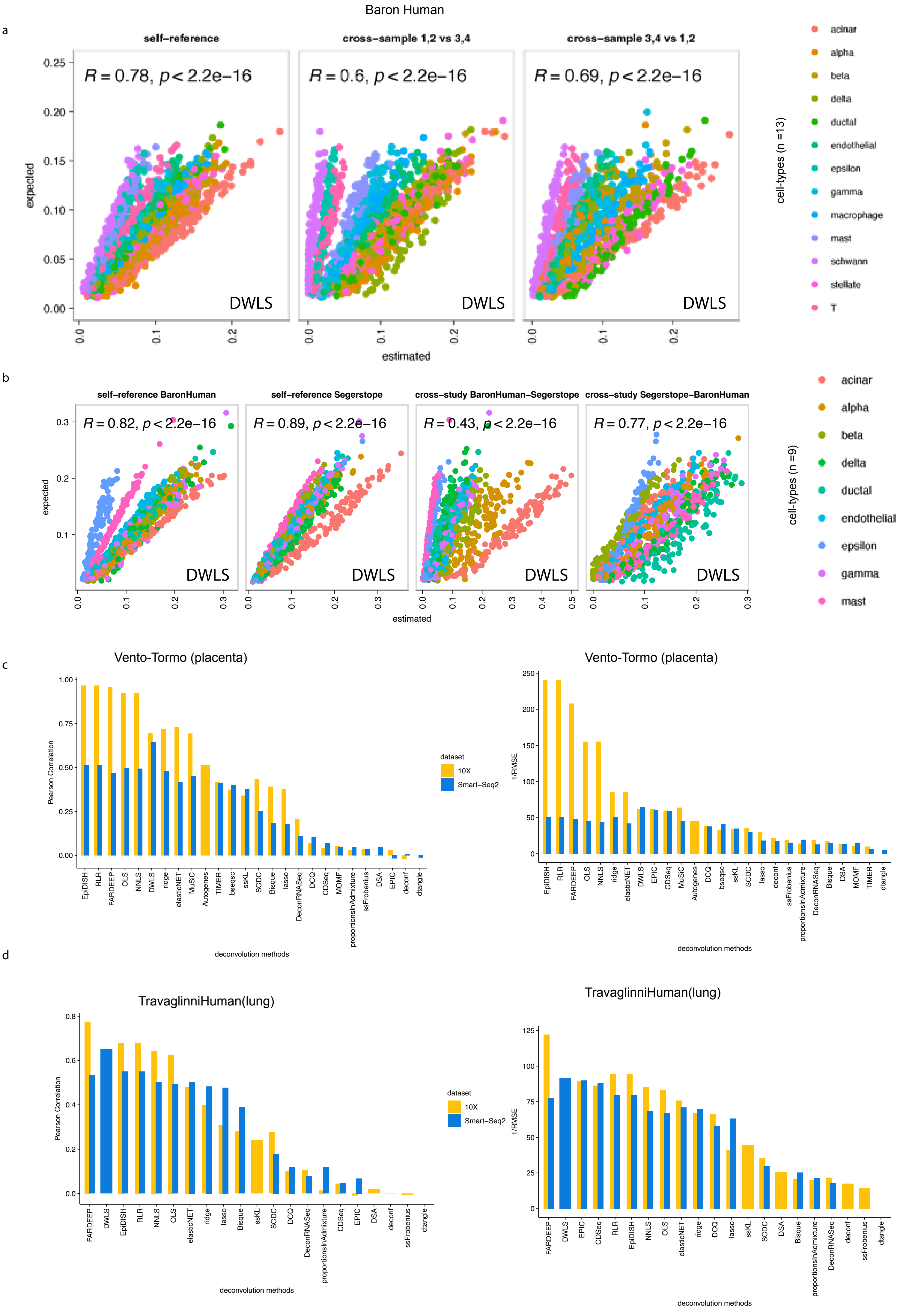

### Supplementary Figure 7

a

KJ Travaglini  2020 ( 10X )

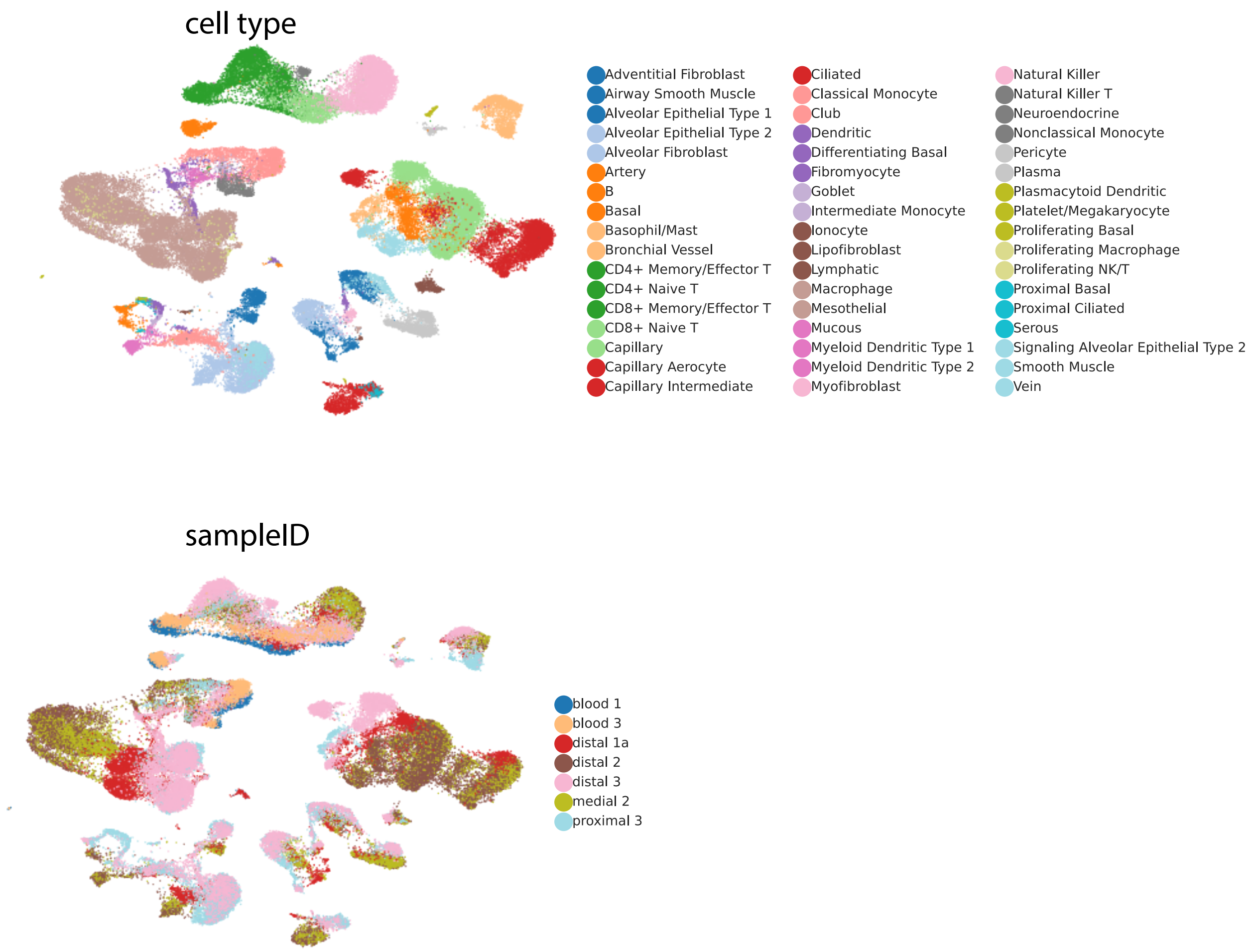


b

KJ Travaglini  2020 -  SS2 analysis of plate-sorted cells

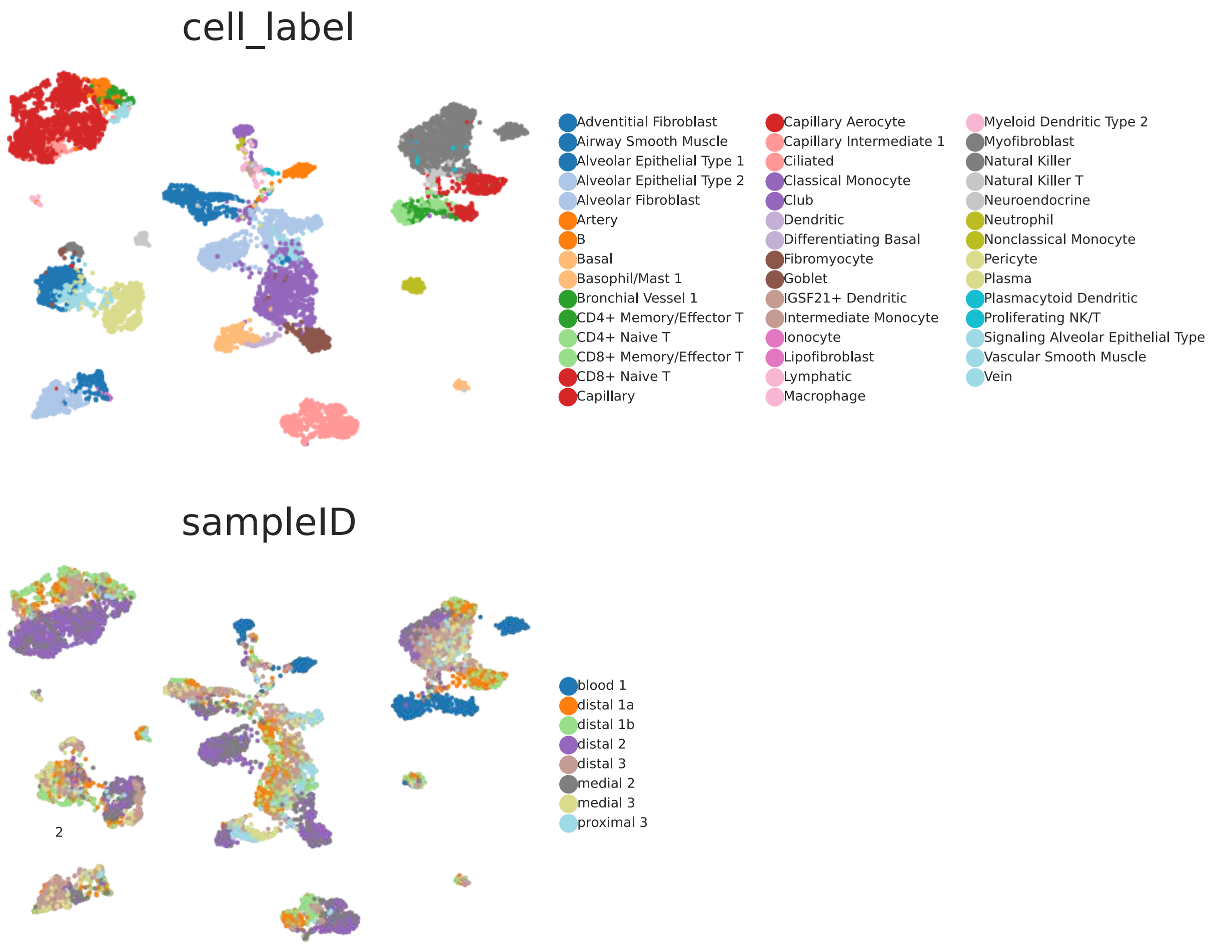

### Supplementary Figure 8

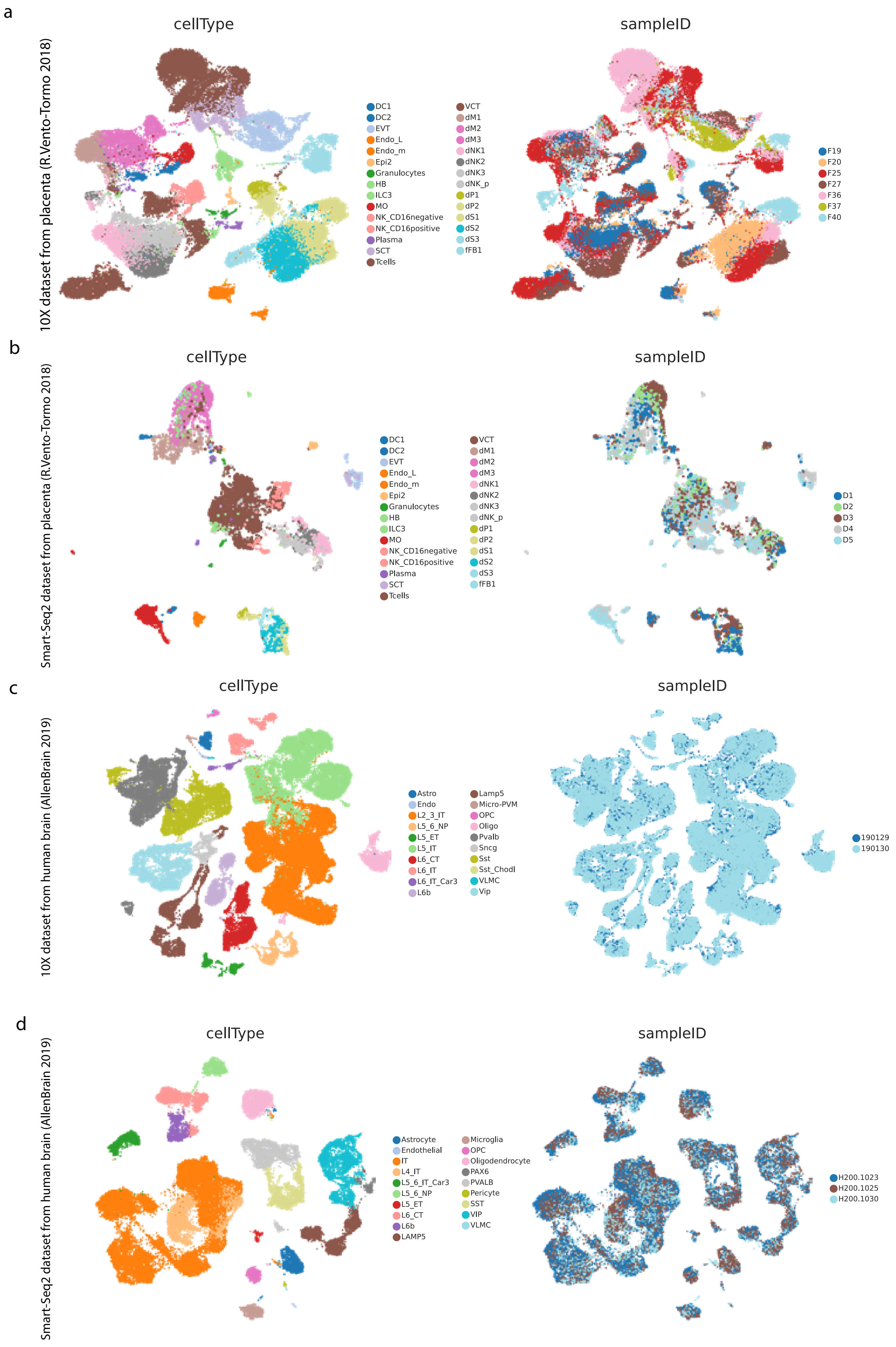

### Supplementary Figure 10

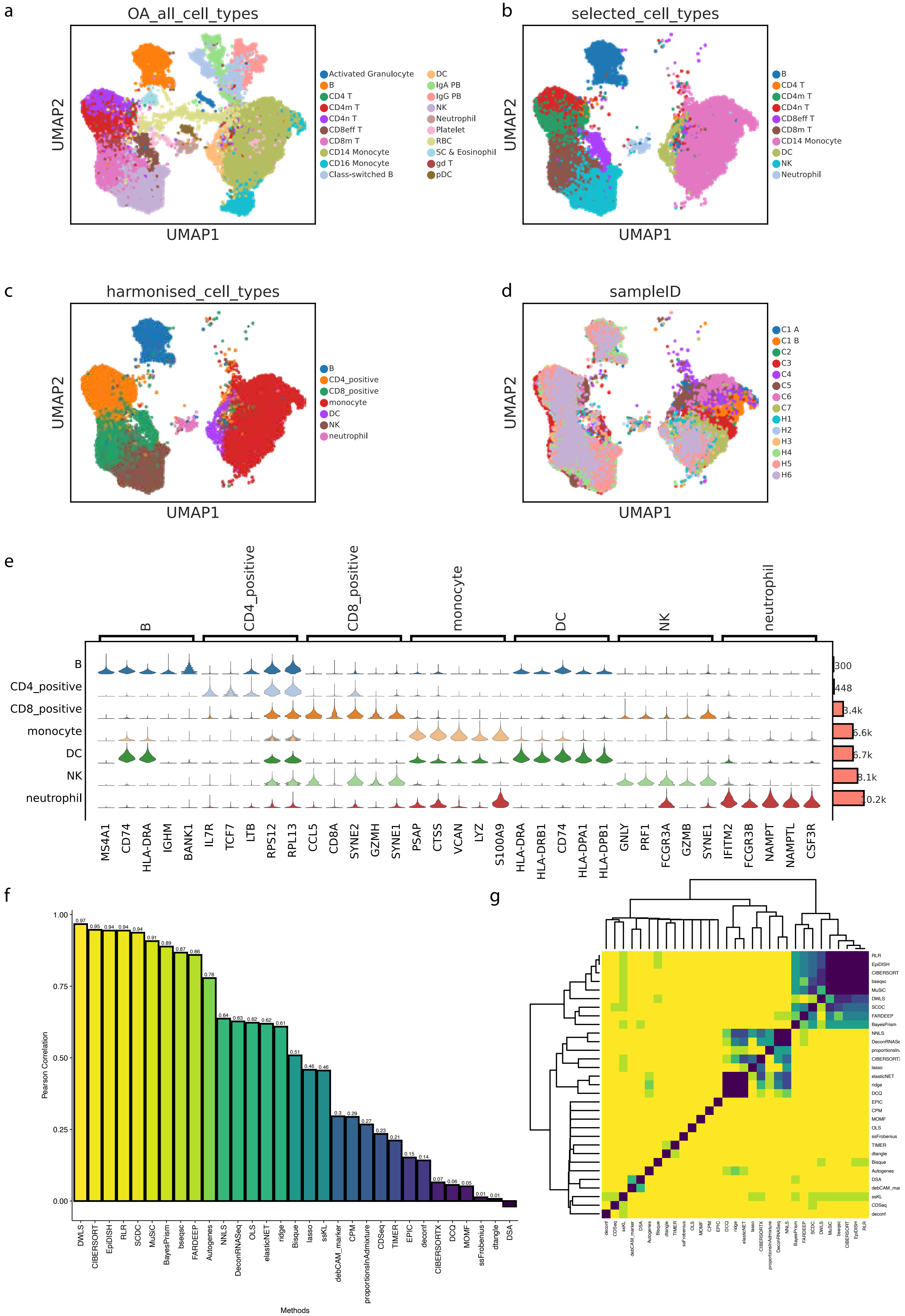

### Supplementary Figure 11

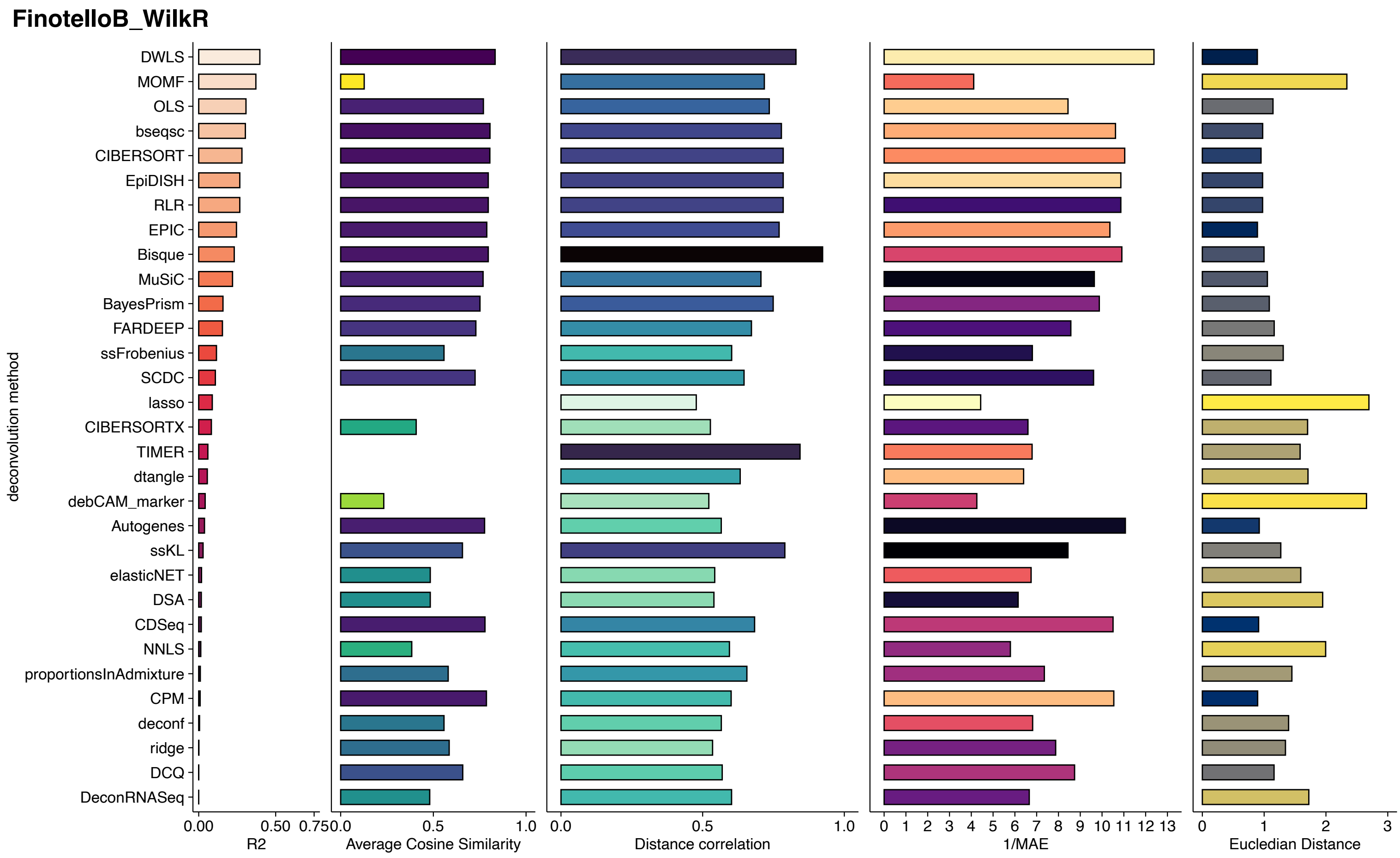

### Supplementary Figure 12

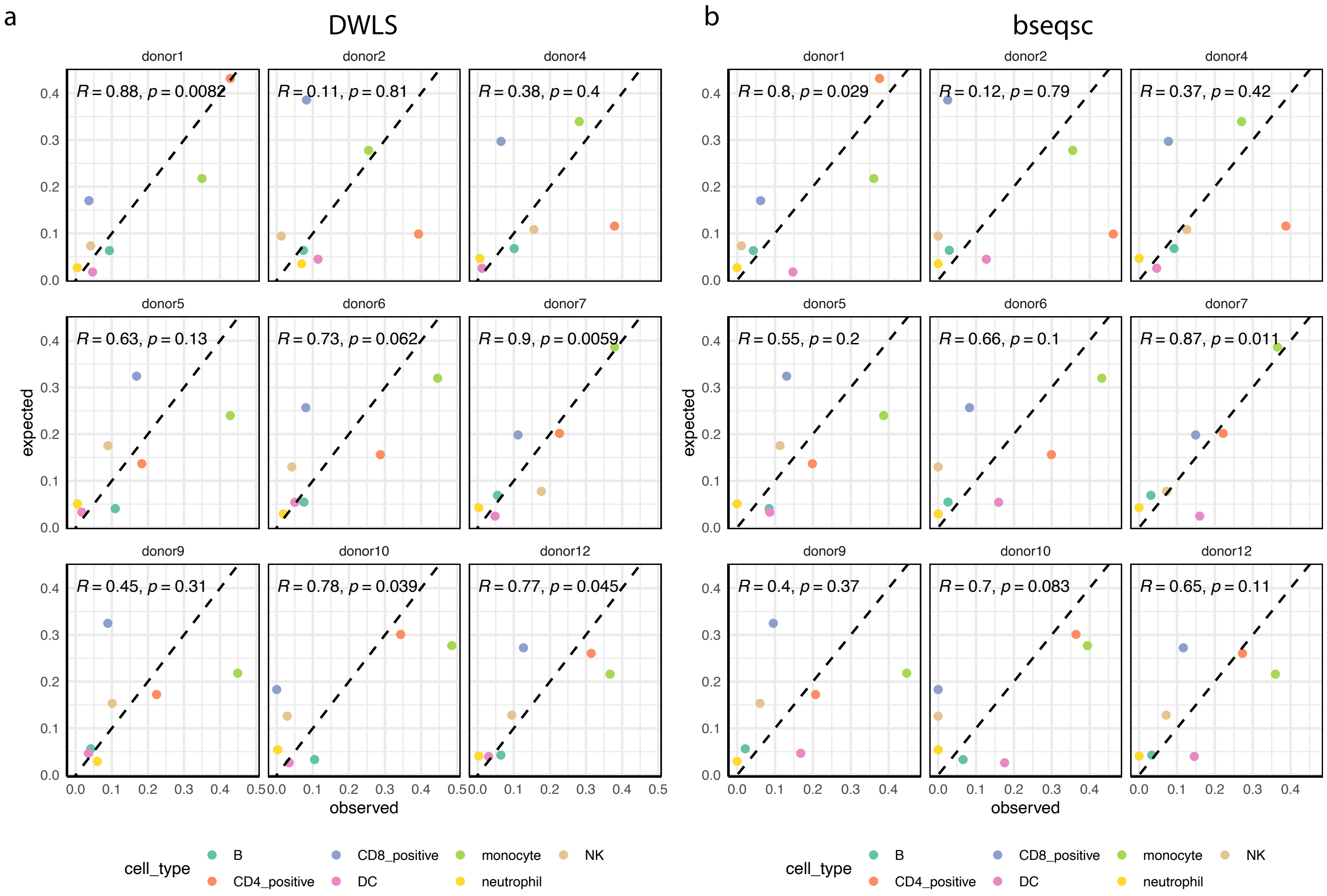

### Supplementary Figure 14

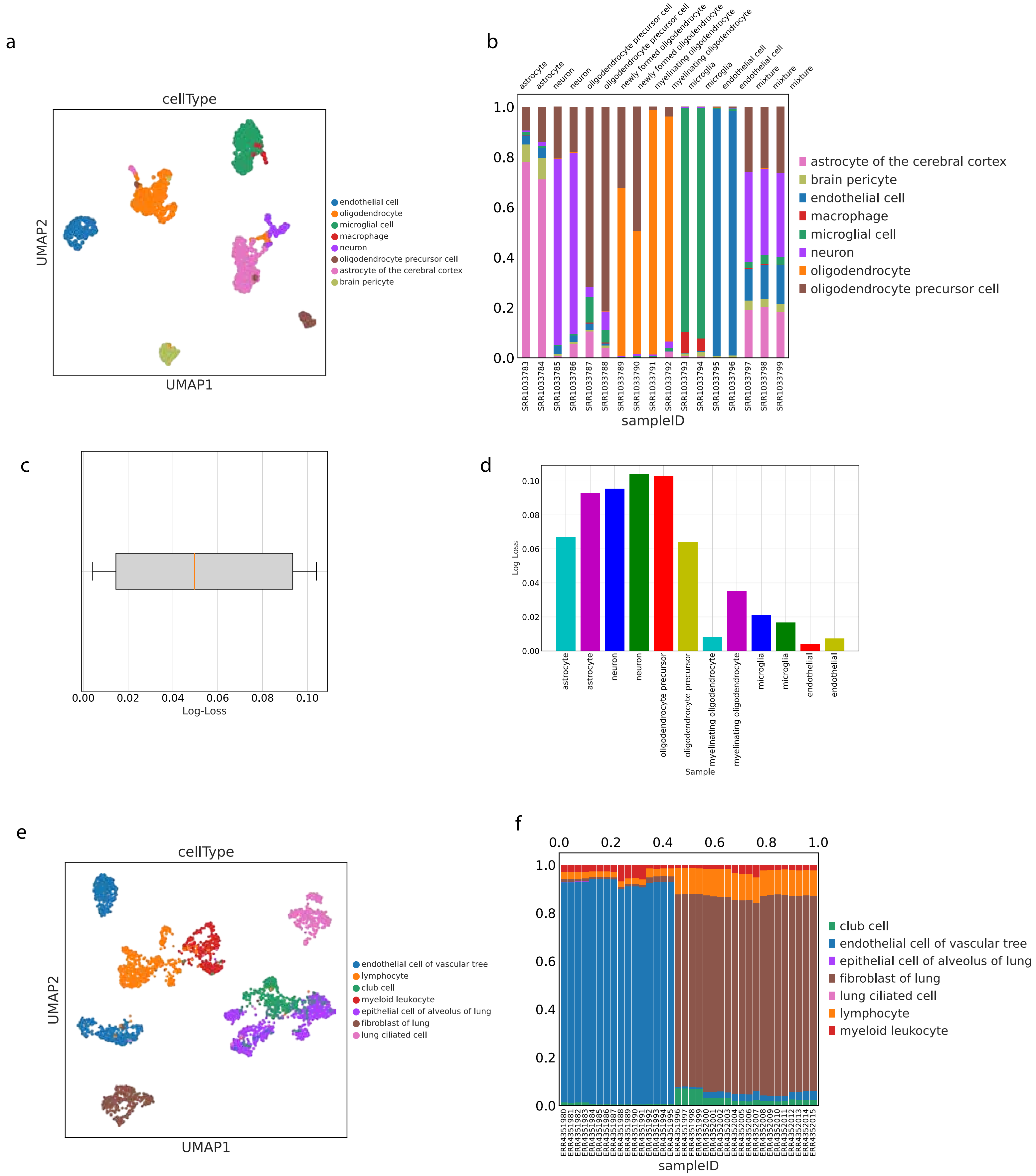

### Supplementary Figure 15

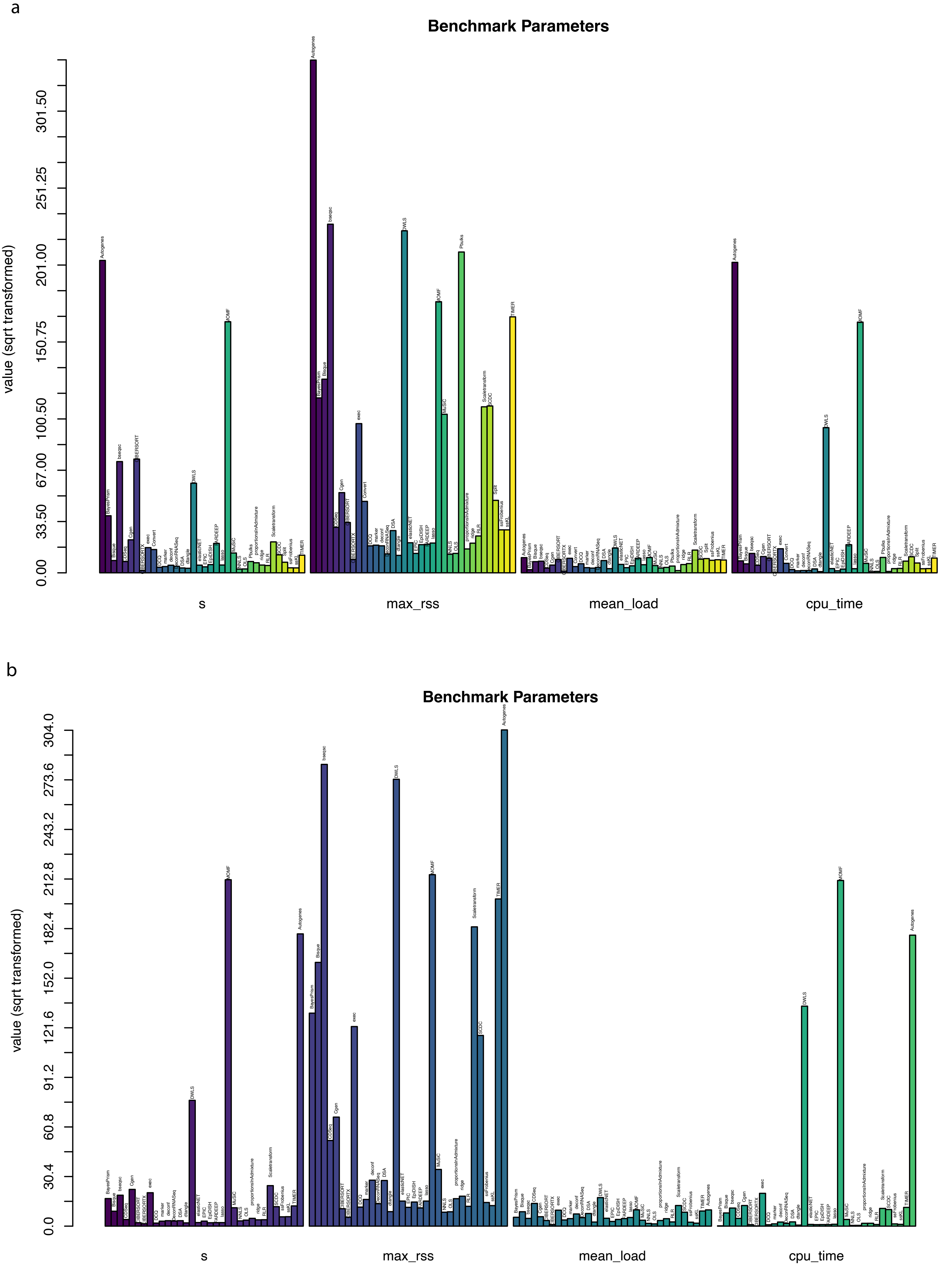

### Supplementary Figure 16

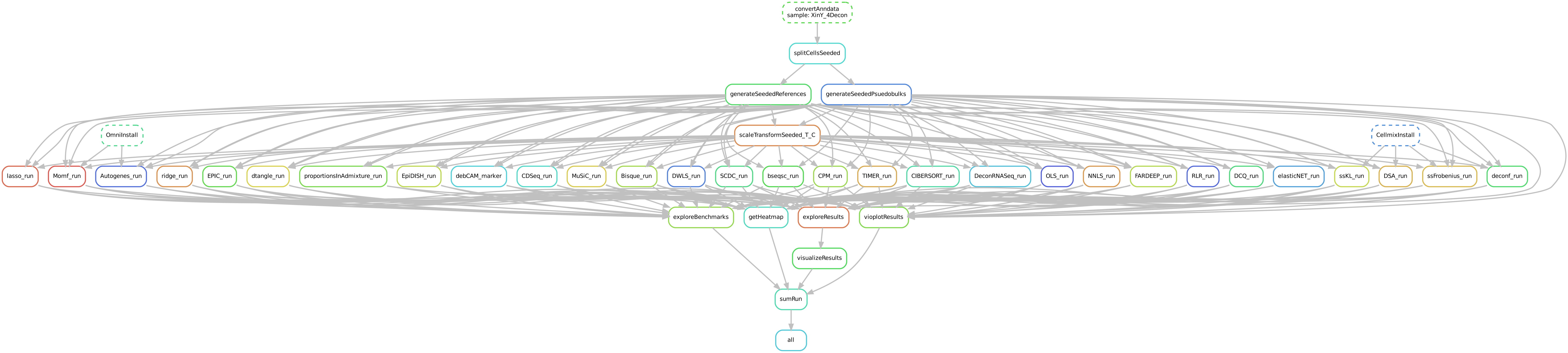

### Supplementary Figure 17

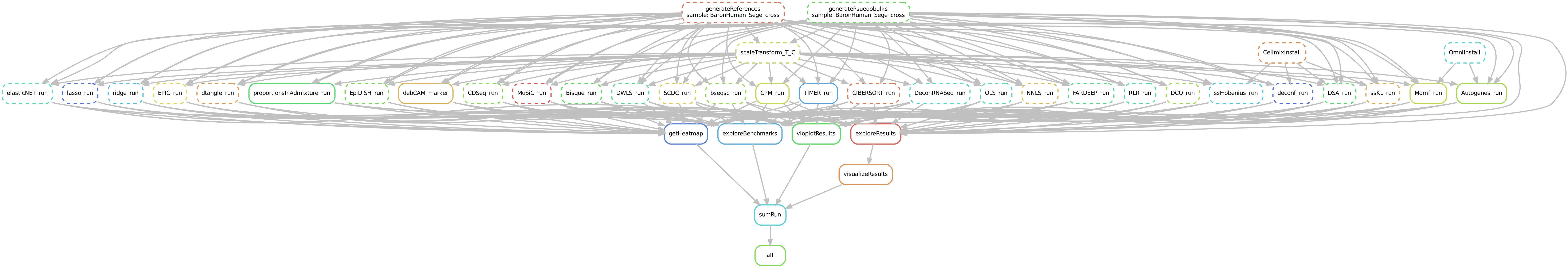

### Supplementary Figure 18

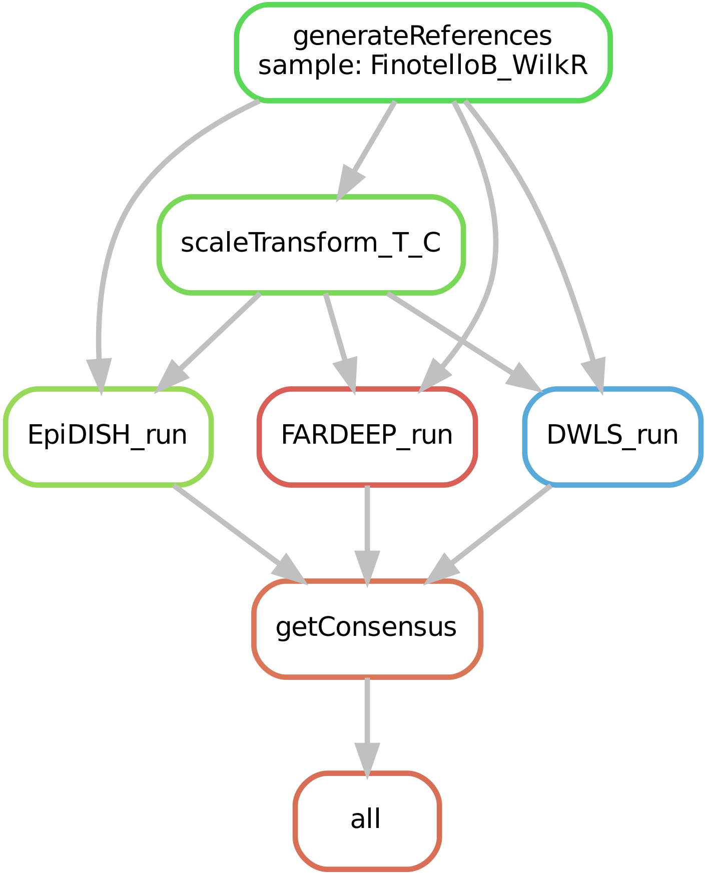
