## Supplementary Figure 9 for "CATD: A reproducible pipeline for selecting cell-type deconvolution methods across tissues"

a

b

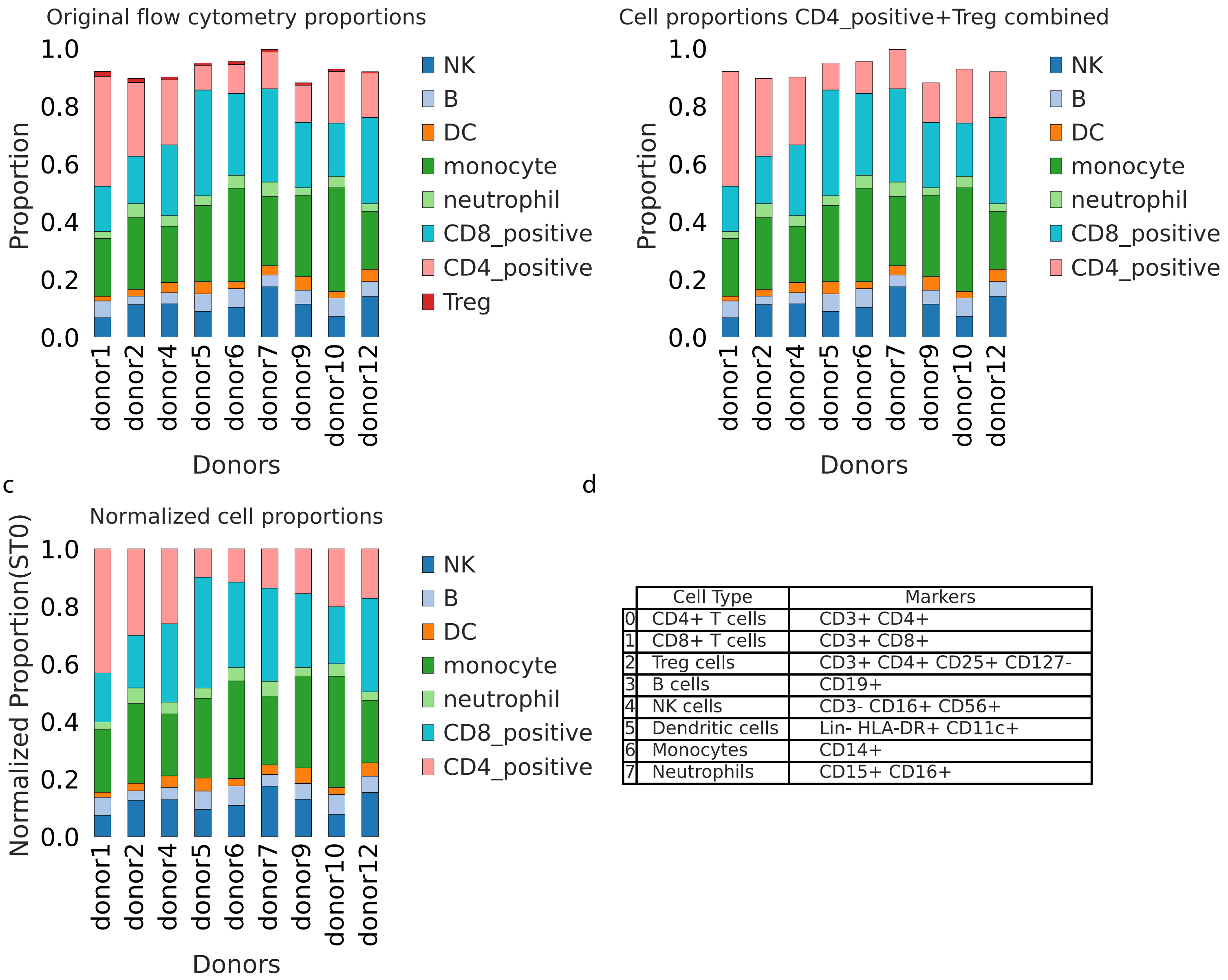


d

| | Cell Type | Markers |
|---|---|---|
| 0 | CD4+ T cells | CD3+ CD4+ |
| 1 | CD8+ T cells | CD3+ CD8+ |
| 2 | Treg cells | CD3+ CD4+ CD25+ CD127- |
| 3 | B cells | CD19+ |
| 4 | NK cells | CD3- CD16+ CD56+ |
| 5 | Dendritic cells | Lin- HLA-DR+ CD11c+ |
| 6 | Monocytes | CD14+ |
| 7 | Neutrophils | CD15+ CD16+ |
