## Supplementary Figure 13 for "CATD: A reproducible pipeline for selecting cell-type deconvolution methods across tissues"

a

FinotelloB_WilkR Reference and Marker−based methods

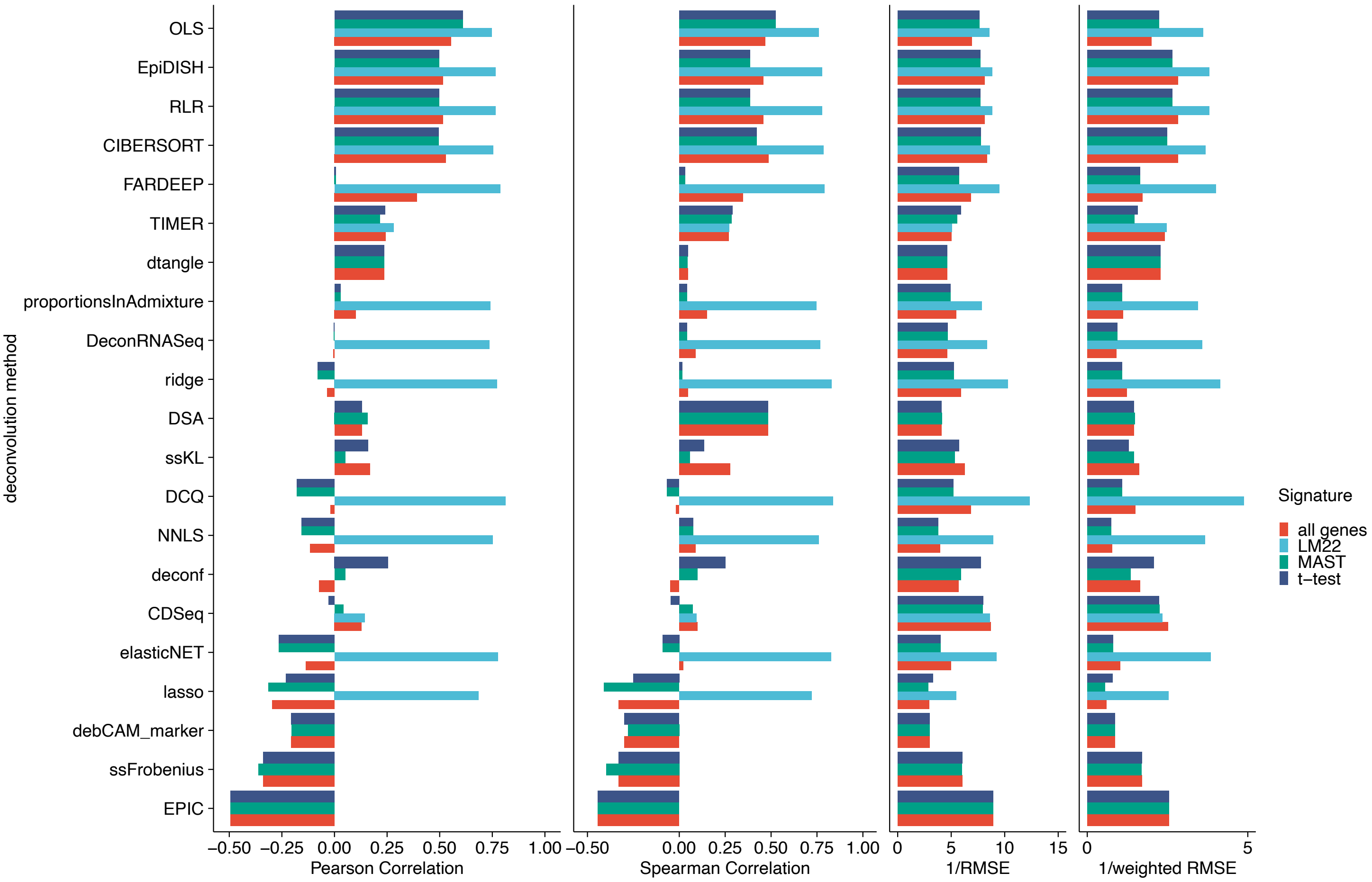


b

FinotelloB_WilkR across different Signatures

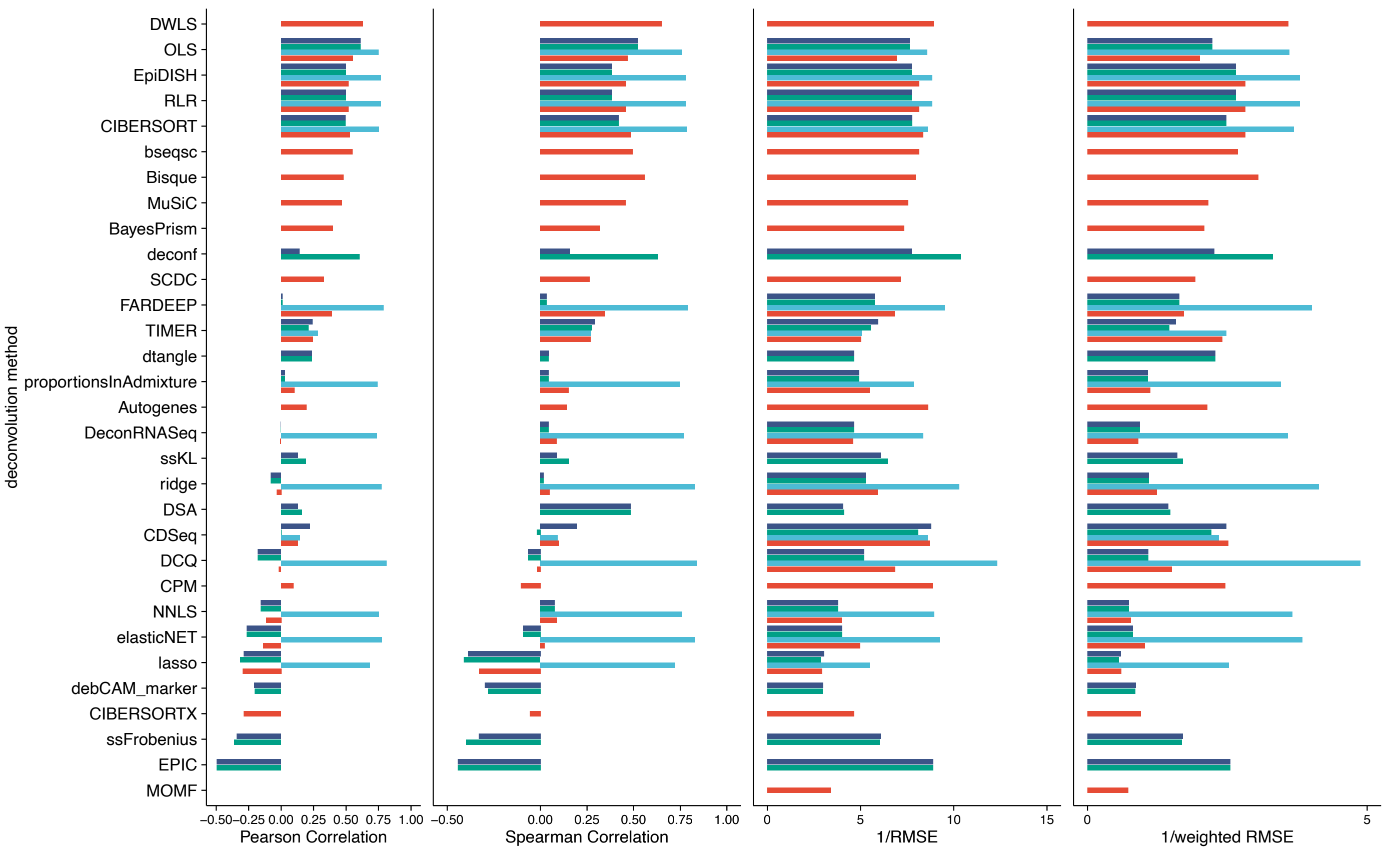
