## Supplementary material for "CATD: A reproducible pipeline for selecting cell-type deconvolution methods across tissues"

**Keywords**

deconvolution, single-cell RNA-seq, benchmarking, cell proportions, pseudo-bulk, real bulk.

**Key Points**

- Thorough assessment of (currently) 31 deconvolution methods, leveraging diverse single-cell RNA-sequencing data from various tissues, alongside extensive simulations and validation against known ground truth data.
- Emphasis on the pivotal role of reference selection, tissue type, and technological nuances in determining the efficacy of deconvolution methods.
- Introduction of the user-friendly and robust Critical Assessment of Transcriptomic Deconvolution (CATD) Snakemake pipeline, enabling efficient and reproducible cell-type deconvolution in real bulk RNA-Seq datasets.

**Abbreviations**

sc-RNA-seq: single-cell RNA-Sequencing

PBMC: Polymorphonuclear/Peripheral Blood Mononuclear cell

PCA: Principal Component Analysis

GEO: Gene Expression Omnibus

GTEX: Genotype-Tissue Expression Project

TCGA: The Cancer Genome Atlas Program

TARGET: Therapeutically Applicable Research to Generate Effective Treatments

**Glossary of key terms**

Cross-reference deconvolution: Supervised deconvolution task where the reference and the pseudo-bulk come from different samples, datasets, or technologies.

Deconvolution :
Cell type deconvolution is a computational technique that aims to infer the relative proportions or abundances of different cell types within a heterogeneous biological sample, such as a tissue or a mixture of cells. This approach leverages gene expression or other molecular data (such as methylation) from a sample to estimate the contributions of individual cell types that make up the complex mixture.

Self-reference deconvolution: Supervised deconvolution task where the reference and the pseudo-bulk come from the same dataset after splitting it into training (reference) and testing (pseudo-bulk).

Snakemake workflow manager:
Snakemake is a workflow management system and domain-specific language used that automates and manages the execution of complex, multi-step data analysis pipelines. It allows users to define the steps, inputs, outputs, and dependencies of a workflow in a human-readable format. Then it automatically manages the execution of these steps, ensuring that they are executed in the correct order while taking advantage of available computational resources efficiently. This tool is particularly valuable in bioinformatics research where reproducibility, scalability, and ease of use are essential.

10X : 10x Genomics' single cell RNA-seq (scRNA-seq) technology, the Chromium Single Cell 3' solution, allows you to analyse transcriptomes on a cell-by-cell basis through the use of microfluidic partitioning to capture single cells and prepare barcoded, next-generation sequencing (NGS) cDNA libraries.

Smart-Seq2(SS2): Smart-seq2 is a single-cell RNA sequencing (scRNA-seq) technology that enables comprehensive transcriptome analysis at the single-cell level. It is characterised by its full-length cDNA synthesis approach, providing high-resolution insights into the gene expression profiles of individual cells.

**Supplementary Methods**

Explanation of the 3 streams of CATD pipeline

1. Self-reference deconvolution pipeline.

Description: For this task we input a single-cell dataset in Seurat or Anndata object format. The dataset is split in two, the training dataset that will be used as a reference and a test dataset that will be used to generate simulated bulk RNA-seq data. The first step of the pipeline includes converting the data from anndata format to seurat object using the sceasy package. Next, the dataset is split in half, the first half will be used for reference generation and the second half for pseudo-bulk generation. In this step the seed can be specified for reproducibility. We also make sure that all the cell types in the single-cell dataset have been included in both halves. The next step of the pipeline happens in parallel and includes the generation of suitable references -that will be later used as inputs in deconvolution methods- and simulated bulk data(see details in the methods below). Once the references and pseudo-bulk files have been generated normalisation and transformation of pseudo-bulk and reference matrices is performed where it is relevant. Next, the relevant generated inputs are fed in the deconvolution methods and cell-type predictions are obtained. The first output of the pipeline is a matrix of cell-type proportions per sample for each deconvolution method which is saved in RDS format under the directory 'Output' . Since in self-reference deconvolution ground truth proportions are known (determined while generating pseudo-bulk profiles) evaluation metrics are calculated across samples for each method. The output of the evaluation is a RDS-formatted file for each metric that includes a list of values of this metric for each deconvolution method. These files are stored under the directory 'Metrics'. Finally, an evaluation of the scalability is performed for each method, the table of values is stored under 'Benchmarks'. The values from the metrics and benchmarks file are used for visualisation of the performance and comparison of scalability across methods. Violin plots and bar charts are used to visualise the above. Resulting plots are then stored under the directory 'Plots'. (Supplementary Figure 11)

2. Cross-reference deconvolution pipeline.

Description: The main steps of the cross-reference pipeline are the same as described in the previous section. However, in this task we need two input single-cell datasets instead of one. The first dataset will be used for the generation of pseudo-bulk(and the ground truth) and the second for the generation of the references. For this task it is required that both datasets originate from the same tissue and hence, they include the same cell-types. In addition, the cell-type labels should be matched to allow evaluation of the results. The primary objective of the cross-reference pipeline is to facilitate the implementation of controlled in silico experiments,through which we can investigate how differences in gene expression between (pseudo)bulk and reference , stemming from sample differences, technologies, or studies, can influence the deconvolution process. Unlike self-reference deconvolution this pipeline introduces more noise and simulates a more realistic deconvolution scenario.(Supplementary Figure 12)

3. Real bulk deconvolution pipeline.

Description: This part of the pipeline was designed for the real bulk deconvolution as well as for future users to deconvolute their own data across the implemented methods. This pipeline is split into two smaller pipelines depending on the presence or absence of ground truth proportions. When ground truth data are present(indicated in the config.yaml file as ***realBulk: 1***) the users need to provide (a) bulk expression matrix to be deconvolved (b) a matrix of known ground truth proportions (c) single-cell data in anndata or Seurat object format. Moreover the users can decide regarding the preprocessing parameters(normalisation transformation of both the bulk and single-cell matrix), which statistical test will be used for marker selection, which deconvolution methods will run, as well as if a consensus of EpiDISH,FARDEEP and DWLS will be calculated. The outputs of the pipeline will be the same as described in the self-reference deconvolution section. In cases where there is no knowledge regarding the ground truth proportions (parameter ***realBulk-noprop : 1*** ) users can still run the pipeline by providing only the bulk and the single-cell reference of their choice. Since in this case evaluation metrics cannot be calculated for each method instead we perform correlation of results across methods to inform the user regarding the concordance of methods in predicting proportions.(Supplementary Figure 13)

**Implementation of deconvolution methods**

Here is the list of deconvolution methods currently implemented in the pipeline ordered alphabetically with their package version numbers.

1. AutogeneS:Implemented from omnideconv package, omnideconv_0.0.0.9000 , autogenes, 1.0.4

2. BisqueRNA :BisqueRNA_1.0.5

3. Bseqsc: bseqsc_1.0

4. CDSeq: CDSeq_1.0.9

5. CIBERSORT: source code available at https://cibersortx.stanford.edu/ upon request. Code was obtained in August 2022.

6. CPM: scBio_0.1.2

7. DCQ : ComICS_1.0.4

8. debCAM_marker : debCAM_1.12.0

9. Deconf : CellMix_1.6.2

10. DeconRNASeq: DeconRNASeq_1.40.0

11. DSA: CellMix_1.6.2

12. dtangle : dtangle_2.0.9

13. DWLS: DWLS_0.1

14. EpiDISH: EpiDISH_2.14.0

15. EPIC: EPIC_1.1.7

16. elasticNET: glmnet_4.1-7

17. FARDEEP: FARDEEP_1.0.1

18. lasso: glmnet_4.1-7

19. MOMF: MOMF_0.2.0, omnideconv_0.0.0.9000

20. MuSiC: MuSiC_1.0.0, TOAST_1.12.0, quadprog_1.5-8, limma_3.54.2, EpiDISH_2.14.1, ggplot2_3.4.2, nnls_1.4

21. NNLS: nnls_1.4

22. OLS: base R, R version 4.2.3 (2023-03-15) lm() function

23. proportionsInAdmixture: energy_1.7-11 WGCNA_1.72-1 fastcluster_1.2.3 dynamicTreeCut_1.63-1 ADAPTS_1.0.22

24. RLR : MASS_7.3-60 energy_1.7-11

25. ridge: glmnet_4.1-7

26. SCDC: SCDC_0.0.0.9000

27. ssFrobenius:CellMix_1.6.2

28. ssKL: CellMix_1.6.2

29. TIMER : This code is adapted from http://cistrome.org/TIMER/download.html. The method is described in Li et al. Genome Biology 2016;17(1):174., PMID 27549193. TIMER Pipeline for analysing immune cell components in the tumour microenvironment. https://github.com/Papatheodorou-Group/CATD_snakemake/tree/main/Modules/TIMER

30.CIBERSORTx , downloaded singularity image, 20 December 2023

31.BayesPrism(omnideconv_0.0.0.9000)

**Implementation of performance evaluation metrics**

The pipeline can evaluate the predictions of deconvolution methods through various metrics when ground truth proportions are available(from pseudo-bulk or real data). The metrics that have been implemented in the pipeline are listed below:

Let:

$X \in R^{NxM}$ be the ground truth matrix, where N is the number of cell types and M is the number of samples

$\widehat{X} \in R^{NxM}$ be the predicted matrix

- N is the number of cell types
- M is the number of samples
- $xij$ represents the element in the $i$-th row in the $j$-th column of matrix $X$
- $\widehat{xij}$ represents the element in the $i$-th row in the $j$-th column of matrix $\widehat{X}$

1. Root Mean Squared error(RMSE)/Root Mean Squared Deviation(RMSD)

$$\text{RMSE/RMSD} = \sqrt{\frac{1}{N \times M} \sum_{i=1}^{N} \sum_{j=1}^{M} (x_{ij} - \widehat{x}_{ij})2}$$

2. Mean Absolute Error(MAE)/Mean Absolute Deviation(MAD)

$$\text{MAE/MAD} = \frac{1}{N\times M}\sum_{i=1}^{N}\sum_{j=1}^{M}\left|x_{ij} - \hat{x}_{ij}\right|$$

3. Pearson Correlation Coefficient(R/pcor)

$$\text{R/pcor} = \frac{\sum_{i=1}^{N}\sum_{j=1}^{M}(x_{ij}-\overline{X}_i)(\hat{x}_{ij}-\overline{\hat{X}}_i)}{\sqrt{\sum_{i=1}^{N}\sum_{j=1}^{M}(x_{ij}-\overline{X}_i)\sum_{i=1}^{N}\sum_{j=1}^{M}(\hat{x}_{ij}-\overline{\hat{X}}_i)2}}$$

Where $\overline{X}_i$ is the mean of the i-th row of the matrix X and $\overline{\hat{X}}_i$ is the mean of the i=th row of matrix $\hat{X}$

4. R-squared

$$R^2 = 1 - \frac{\sum_{i=1}^{N}\sum_{j=1}^{M}(x_{ij}-\hat{x}_{ij})2}{\sum_{i=1}^{N}\sum_{j=1}^{M}(x_{ij}-\overline{X}_i)2}$$

5. Spearman Correlation Coefficient(ρ/rho)

$$\rho = 1 - \frac{6\sum_{i=1}^{N}\sum_{j=1}^{M}d_{ij}^2}{N(N^2-1)\cdot M}$$

where $d_{ij}$ = R($X_{ij}$) - R($\hat{X}_{ij}$) is the difference between the ranks for the element in the i-th row and the j-th column for each pair of observations.

6. Cosine Similarity(cos)

$$\cos(x, \widehat{x}) = \frac{\sum_{i=1}^{N}\sum_{j=1}^{M} x_{ij} \cdot \widehat{x}_{ij}}{\sqrt{\sum_{i=1}^{N}\sum_{j=1}^{M} (x_{ij})2} \cdot \sqrt{\sum_{i=1}^{N}\sum_{j=1}^{M} (\widehat{x}_{ij})2}}$$

7. Distance correlation

Let $x$ be a vector of values from ground truth matrix X and $\widehat{x}$ be a vector of values from predicted matrix $\widehat{X}$

$$dCov(x, \widehat{x}) = \frac{1}{n^2} \sum_{i=1}^{n}\sum_{j=1}^{n} D(x_i, x_j)\, D(\widehat{x}_i, \widehat{x}_j)$$

$$dCor(x, \widehat{x}) = \frac{dCov(x, \widehat{x})}{\sqrt{dCov(x,x)\, dCov(\widehat{x}, \widehat{x})}}$$

dCov & dCor = Distance covariance & correlation as described in this study[50]

Note that dCov(X,X) = dVar(x), which is the distance variance.

n = number of observations(number of values in each vector $x$ and $\widehat{x}$

$D(x_i, x_j)$ = centered Euclidean distances between the i_th and j_th observations of x

$D(\widehat{x}_i, \widehat{x}_j)$ = centered Euclidean distances between the i_th and j_th observations of y

8. Minkowski distance

$$\text{Minkowski Distance} = \left(\sum_{i=1}^{N}\sum_{j=1}^{M} |x_{ij} - \widehat{x}_{ij}|^{p}\right)^{1/p}$$

For p = 2 it is simplified to **Euclidean Distance** which is the metric used in the CATD pipeline.

9. Weighted RMSE

Let $w_i$ be the weight assigned to the i-th cell type based on its ground truth proportion. The weighted RMSE(wRMSE) can then be calculated as:

$$\text{RMSE/RMSD} = \sqrt{\frac{1}{N\times M}\sum_{i=1}^{N}\sum_{j=1}^{M} w_i(x_{ij} - \hat{x}_{ij})2}$$

$w_i$ is defined as the inverse of the ground truth proportion for the i-th cell type

$w_i = 1/p_i$

Where $p_i$ is the ground truth proportion for the i-th cell type.

This modification of RMSE ensures that rare cell types, characterised by lower ground truth proportions, contribute more significantly to the weighted RMSE.

All the metrics have been estimated in R and the functions can be found here https://github.com/Functional-Genomics/CATD_snakemake/blob/main/Modules/Res_explore/Res_explore.R. For each metric, the scores of the methods are saved as named list objects throughout the pipeline. Graphs are also constructed from these lists for initial viewing of the results.

**Supplementary Figure 1. Multi-Parametric simulation for pseudo-bulk RNA-seq data deriving from single-cell RNA-Seq data** a,b UMAP plots of 10x mouse brain single-cell dataset after filtering and reanalysis. The dataset contains 16 cell-types as shown with different colours (left) and 28 samples. c. bar plot of cell-type proportions for each sample based on the single-cell dataset. c. violin plot of the cosine similarity of gene expression in randomly sampled pseudo-bulks with 100,500 and 1000 samples (left) as well as across different numbers of sampled cells 100, 10K and 100K (right). d. Violin plots showing the distribution of cell fractions across different sampling methods in 16 cell types from Hrvatin.

**Supplementary Figure 2. Comparison of sampling methods in terms of variance and gene correlation.**

**a**. Scatter plot comparing the coefficient of variation between bulk and pseudo-bulk profiles across methods adapted from code by Hu and Chikina. b. Heatmaps demonstrating the pairwise gene correlation across simulation methods and in comparison with real data.

**Supplementary Figure 3: Preprocessing of simulated data from 6 brain tissue datasets.** a-f. Barplots showing Pearson Correlation values of 6 self-reference deconvolution tasks(using 6 single-cell datasets from brain tissue) in which the effect of scaling or transforming first the input matrices is tested.

**Supplementary Figure 4 Comparison of gene expression in pseudo-bulk (sc) and real bulk data.**

a.UMAP plot of a single-cell dataset (BaronHuman2018) from 4 human pancreatic islets after re-analysis,different colours indicate different cell-types based on author's annotations (left) or the different samples(right). b.scatter plot showing the differences in expression between summarised single-cell (BaronHuman2018) and a sample from bulk dataset (Fadista2015) shown in blue colour, as well as real bulk versus real bulk profiles (purple) and pseudo-bulk versus pseudo-bulk profiles(green) c. Scatter plot demonstrating gene expression between 12 different bulk samples from Fadista2015 pancreatic RNA-seq dataset. d. scatter plot of gene expression deriving from 12 pseudo-bulk profiles generated from the pancreatic single-cell dataset

**Supplementary Figure 5. Combined Principal Component Analysis of real bulk and pseudo-bulk samples.**

a.PCA results from 78 real bulk samples (Fadista 2014) and 78 pseudo-bulk samples (BaronHuman2018). PC1 and PC2 can explain 80.46% of the variance (left) , PC2 and PC3 shown on the right scatter plot. b. Variance ratio plot, "Explained" variance ratio as a function of the number of principal components. The inclusion of 30 PCs accounts for 98.08% of the variance.

**Supplementary Figure 6. Cross-reference deconvolution of pseudo-bulk samples across different tissues.**

a. scatter plots showing results from self-reference and 2 examples of cross-sample deconvolution from the method DWLS utilises a single-cell from human pancreas (13 cell-types). Pearson correlation (R) and p-value are reported for each task. b.scatter plots showing the results from two self-reference tasks from two pancreatic studies (9 cell-types) as well as two cross-study deconvolution tasks. c. Bar plots showing 1/RMSE and Pearson correlation values from the 26 feasible deconvolution runs from two placenta tissue datasets that come from the same study but different technologies (10X and Smart-seq2). d. Bar plots showing 1/RMSE and Pearson correlation values from the 26 feasible deconvolution runs from two lung tissue datasets that come from the same study but different technologies (10X and Smart-seq2)

**Supplementary Figure 7. Single-cell references from human lung tissue.**

a.UMAP plots of a single-cell 10X dataset from human lung after re-analysis, different colours indicate different cell-types based on author's annotations (top) or the different samples (bottom). b. UMAP plots of a single-cell Smart-seq2 dataset from human lung after re-analysis, different colours indicate different cell-types based on author's annotations (top) or the different samples (bottom).

**Supplementary Figure 8.Single-cell references from human placenta and brain for cross-technology deconvolution**

a.UMAP plots of two single-cell dataset(10X and Smart-Seq2) from human placenta after re-analysis, different colours indicate different cell-types based on author's annotations (left) or the different samples (right). b. UMAP plots of two single-cell dataset(10X and Smart-Seq2) from human brain after re-analysis, different colours indicate different cell-types based on author's annotations (left) or the different samples (right).

**Supplementary Figure 9. Flow cytometry measurements from real PBMC data**

Barplots showing the cell type fractions across samples/donors

A.original experimental proportions b. after summarising CD4 positive and T-regulatory cells c. after normalising data from b to sum to one. d. cell type markers that were used for the flow cytometry measurements.

**Supplementary Figure 10. Single-cell reference for deconvolution.**

UMAP representations of single-cell reference from Wilk et al. a. Original authors annotations b. Selected cell types to map to bulk profiles. C. harmonised names to match cell proportions from bulk PBMC data. d. demonstration of sampleIDs across cell types. e. Top 5 marker genes per cell type after Differential expression analysis across cell type to confirm the authors' annotation. f. Barplot showing deconvolution results across methods in the self-reference task for the single-cell PBMC data set. g. Correlation plot showing the agreement between deconvolution results in the PBMC self-reference task.

**Supplementary Figure 11. Evaluation of deconvolution methods in the real bulk deconvolution task across 5 additional metrics**

**Supplementary Figure 12 Deconvolution results of real PBMC data per donor**

a. scatter plots showing the 'per sample/donor' deconvolution results (DWLS method) of the real PBMC dataset. Different R values are observed across samples. d. Same results for the bseq-sc deconvolution method.

**Supplementary Figure 13. Evaluating deconvolution methods across different signatures,**

a. Barplot shows the performance of methods across 4 different cases (all genes, LM22 signature, MAST-DE genes, DE genes t-test) in reference and marker based approaches b. Summarised results across all methods.

**Supplementary Figure 14. Application of consensus deconvolution approach in samples of purified cell types**

a. UMAP plot of downsampled single-cell RNA-seq dataset from TabulaMuris (cerebral cortex subset) composed of 8 cell-types. Original authors' labels have been reduced(Methods). b. Bar plot showing the proportions of cell-types for each sample in the bulk mouse cerebral cortex dataset as predicted by the consensus deconvolution approach. c. boxplot illustrating the distribution of log-loss values across samples when comparing the predicted cell types from deconvolution with the known presence of a cell type in a purified sample. d. The bar plot shows the log-loss values for 12 samples in the dataset (SRR1033789, SRR1033789 -newly formed oligodendrocytes and the mixtures- SRR1033797, SRR1033798, SRR1033799 were excluded from this analysis). Lower the log-loss indicates more accurate predictions of the method. e. UMAP plot of the downsampled single-cell RNA-seq dataset from the lung subset of GTEx composed of 7 cell-types. Original authors' labels have been reduced (Methods). f. Barplot showing the proportions of cell-types for each sample in the bulk human fetal lung dataset[known purified endothelial (left) and non endothelial (right) samples] as predicted by the consensus deconvolution approach.

**Supplementary Figure 15. Evaluation of scalability on simulated and real PBMC data.**

a-b. for scalability assessment purposes, time (s), memory (maximum resident set size), mean load (CPU usage divided by total time) and CPU time are visualised in barplot for Wilk2020-self-reference task as well as for the deconvolution of Finotello PBMC real bulk samples.

**Supplementary Figure 16. Directed acyclic graph(DAG) for self-reference task of the CATD pipeline.**

**Supplementary Figure 17. Directed acyclic graph(DAG) for cross-reference task of the CATD pipeline.**

**Supplementary Figure 18. Directed acyclic graph (DAG) for real bulk consensus deconvolution of the CATD pipeline.**
